# Sex differences in electrophysiological properties of mouse mPOA neurons revealed by *in vitro* whole-cell recordings

**DOI:** 10.1101/808592

**Authors:** Wen Zhang, Shuai-shuai Li, Zhuo-lei Jiao, Ying Han, Zi-yue Wang, Xiao-Hong Xu

## Abstract

The medial preoptic area (mPOA) of the hypothalamus is sexually dimorphic and controls sexually dimorphic display of male mating and parental care. Yet, despite extensive characterization of sex differences in the mPOA, we know surprisingly little about whether or how male and female mPOA neurons differ electrophysiologically, which relate more directly to neuronal firing and behavioral pattern generation. In this study, we performed whole-cell patch clamp recordings of the mPOA in acute brain slices cut from virgin adult mice, and compared in total 29 electrophysiological parameters between male and female mPOA neurons. We find that resting membrane potential (*V*_m_), input resistance (*R*_m_), time constant (*τ*_m_), threshold (*V_th_*) and minimum current (rheobase) required to generate an action potential differ significantly between male and female in a cell-type dependent manner. Nonetheless, there is little evidence for profuse sex differences in neuronal excitability, except for a higher probability of rebound neurons in males. Depletion of male gonadal hormones in adulthood partially de-masculinizes sexually dimorphic electrophysiological parameters, suggesting that some of these sex differences may establish during development. Furthermore, as a demonstration of the behavioral relevance of these sex differences, we show that pharmacologic blockage of currents mediated by T-type Ca^2+^ channel, which underlie rebound and tends to be larger in male mPOA neurons, result in behavioral deficits in male mating. In summary, we have identified key sex differences in electrophysiological properties of mPOA neurons that likely contribute to sexually dimorphic display of behaviors.

**Significance Statement:** Sex represents an important biological variable that impact an individual’s behaviors, physiology and disease susceptibility. Indeed, sex differences in the nervous system manifest across many different levels and scales. Yet, throughout previous multifaceted investigations on sex differences in the brain, electrophysiological characterizations, which could potentially bridge cellular and molecular sex differences with sexually dimorphic brain functions and behaviors, remains scant. Here, focusing on an evolutionarily conserved sexually dimorphic nucleus, we investigated sex differences in electrophysiological properties of mPOA neurons and its modulation by gonadal hormones in adult males via *in vitro* whole-cell patch clamp. As a result, we identified novel sex differences in electrophysiological properties that likely contribute to sexually dimorphic display of behaviors and physiological functions.

## Introduction

The medial preoptic area (mPOA) of the hypothalamus functions as an essential node in an evolutionarily conserved neural network that controls social behaviors (Newman, 1999, O’Connell and Hofmann, 2012, Zha and Xu, 2015). Notably, it is sexually dimorphic in nearly all species examined to date (Gorski et al., 1978, Panzica et al., 1996, Orikasa and Sakuma, 2010, Campi et al., 2013), including humans (Swaab and Fliers, 1985, Hofman and Swaab, 1989). Sexually dimorphic features of the mPOA include neuronal numbers/densities (Gorski et al., 1978, Sickel and McCarthy, 2000), synaptic organization (Raisman and Field, 1973, Ayoub et al., 1983), distributions of innervating fibers (Simerly et al., 1984) and gene expression levels (Xu et al., 2012). These features are sensitive to hormonal manipulations during development and in adulthood (Raisman and Field, 1973, Orikasa and Sakuma, 2010, Xu et al., 2012). Functionally, sexually dimorphic activation of mPOA neurons underlies sexually dimorphic display of male mating and pup care in mice (Wu et al., 2014, Wei et al., 2018).

Despite extensive characterization of the mPOA in the literature, it remains largely unknown whether or how male and female mPOA neurons differ electrophysiologically. As electrophysiological properties reflect the distributions of voltage-dependent ion channels (Brumberg et al., 2000, Tripathy et al., 2017, Zhang et al., 2019), it is perceivable that sex differences in the expression of ion channels or synaptic organization may result in sexually dimorphic electrophysiological features of mPOA neurons that contribute directly to sex-specific activation patterns underlying sexually dimorphic display of behaviors. For instance, more depolarized resting membrane potential together with lower action potential threshold could in theory lead to easier firing of some neurons (Platkiewicz and Brette, 2010). Similarly, distinctive firing modes such as regular or bursting firing could encode different motivational states to regulate behaviors (Pan et al., 2016, Stagkourakis et al., 2018, Yang et al., 2018). In fact, sex differences in neural excitability have been identified for striatal medium spiny neurons in rats (Dorris et al., 2015, Cao et al., 2018). Additionally, gonadal hormone sensitive electrophysiological properties have also been documented for neurons in the anteroventral periventricular nucleus (AVPV) in mice (Piet et al., 2013, Wang et al., 2016) and for neurons in the song production pathway in birds (Roberts et al., 2007; Liu et al., 2010). Regrettably, while electrophysiological characterization of the mPOA have been performed previously (Hodgkiss and Kelly, 1990, Hoffman et al., 1994), these few studies did not include female subjects for comparison of sex differences. Nor did they investigate hormonal regulation of mPOA electrophysiological properties.

We reason that understanding sex differences in electrophysiological properties of the mPOA will provide a bridge to connect known anatomical and molecular sex differences with sex-specific display of behaviors and will facilitate understanding of neural mechanisms that govern sexually dimorphic brain functions. To this end, we obtained electrophysiological measurements of mPOA neurons in acute brain slices cut from virgin adult male and female mice of a wildtype strain or of a transgenic mouse line that specifically labels Vglut2+ (vesicular glutamate transporter 2) neurons. In addition, we compared electrophysiological properties of mPOA neurons in castrated males compared to sham controls. As the result, we find that several key electrophysiological parameters including both passive and active membrane properties differ significantly between male and female in a cell-type dependent manner but there is little evidence for profound sex differences in neuronal excitability, except for a higher probability of rebound neurons in male. Furthermore, depletion of male gonadal hormones in adult only partially de-masculinizes/feminizes electrophysiological parameters of mPOA neurons. Finally, as a demonstration of the behavioral relevance of these electrophysiological sex differences, we show that pharmacologic blockage of T-type Ca^2+^ channels, which mediate the rebound current, diminishes male mating behaviors. Together, our study uncovers novel sex differences in electrophysiological properties of mPOA neurons that likely subserve sex differences in behaviors and/or other physiological functions.

## Materials and Methods

### Animals

Adult C57BL/6 male and female mice were purchased from the Slac Laboratory Animal (Shanghai). *Slc17a6^tm2(cre)/Lowl^*/J (referred to as *Vglut2-Ires-Cre*; Cat # 016963) and B6;129S6-*Gt(ROSA)26Sor^tm9(CAG-tdTomato)Hze^*/J (referred to as *Ai9*; Cat # 007905) were purchased from the Jackson Laboratory and bred in house. Male and female mice were group housed (5 animals maximum/per cage) according to the sex. Castration and sham surgery were performed as previously described (Xu et al., 2012; Wei et al., 2018) with animals anesthetized under intraperitoneal (i.p.) injection of ketamine (80 mg/kg) and xylazine (8 mg/kg). Animals were allowed 3 to 4 weeks to recover after surgeries before sacrifice for *in vitro* electrophysiological recording. All animals were housed in the Institute of Neuroscience animal facility on 12h light/dark cycle with food and water *ad libitum* until use. All animals were aged between 8 to 22 weeks at the time of sacrifice and were age-matched for comparison between groups. All experimental protocols were approved by the Animal Care and Use Committee (IACUC No. NA-016-2016).

### Preparation of acute brain slices

Adult mice were anesthetized with isoflurane, perfused transcardially with ice-cold oxygenated (95% O_2_/5% CO_2_) high-sucrose solution (in mM: 2.5 KCl, 1.25 NaH_2_PO_4_, 2 Na_2_HPO_4_, 2 MgSO_4_, 213 sucrose, 26 NaHCO_3_), and decapitated. Afterwards, the brain was quickly extracted into ice-cold oxygenated high-sucrose solution and sectioned in the coronal plane at 250 μm in the same buffer on a vibratome (VT1200S; Leica). After sectioning, slices containing mPOA were identified and incubated in oxygenated artificial cerebrospinal fluid (ACSF; in mM: 126 NaCl, 2.5 KCl, 1.25 NaH_2_PO_4_, 1.25 Na_2_HPO_4_, 2 MgSO_4_, 10 Glucose, 26 NaHCO_3_, 2 CaCl_2_) at 34°C for 1h.

### Hormone assays

Trunk blood was collected at the time of killing before perfusion. Serum was prepared by collecting the supernatant of blood after 7500 *g* centrifugation in 4 °C. Hormone titers were assayed using a testosterone ELISA kit (DRG Instruments GmbH, Germany, Division of ARG international, Inc, Cat# EIA-1559) according to the manufacturer’s protocols.

### Whole-cell patch recordings

Slices were transferred to a recording chamber, superfused at 1-2 ml / min with oxygenated ACSF, and held in position by nylon fibers glued onto a U-shaped platinum frame. Neurons were visualized with an upright, infrared-differential interference contrast microscope (BX51WI; Olympus) that is equipped with an infrared CCD camera (IR-1000; DAGE-MTI). Patch pipettes were ∼2 μm in diameter at the tip and had impedances of 3-5 MΩ when filled with intracellular solution (in mM: 135 K-gluconate, 4 KCl, 10 HEPES, 10 sodium phosphocreatine, 4 Mg-ATP, 0.3 Na_3_-GTP and 0.5 biocytin; pH:7.2; 265 mOsm).

All recordings were acquired using Multi-clamp 700B amplifier (Molecular Devices) and Digi-data 1440A interface (Molecular Devices). To assess the intrinsic membrane properties, neurons were hold at −60 mV under current clamp, and injected with positive current steps (1000 ms in duration) and negative current steps (800 ms in duration) from 0 until firing saturation (at least 100 pA) in 10 pA increments and 20 s intervals. Repeated 20 pA negative current steps (500 ms in duration) were delivered at the end of each sweep to obtain the passive membrane properties. Afterwards, spontaneous postsynaptic currents were recorded under voltage clamp mode with membrane potential holding at −70 mV for EPSCs recording and 0 mV for IPSCs recording.

To record voltage-dependent Ca^2+^ currents, an online P/4 protocol was applied to minimize leak and residual capacitive currents. During the recording, tetrodotoxin (TTX; 1.5 μM, Absin), 4-aminopyridine (4-AP; 5 mM, Alomone Labs) and XE-991 (10 μM, Sigma-Aldrich) was added to the bath solution and tetraethylammonium (TEA; 2 mM, Sigma-Aldrich) was added to the internal solution to block Na^+^ and K^+^ channels. Low-voltage-activated Ca^2+^ channels (T-type Ca^2+^ channels) were activated by a −35 mV test pulse (500 ms) that is preceded by a −100 mV pre-pulse (200 ms). The currents mediated by T-type Ca^2+^ channels were extracted by subtracting current recorded after application of mibefradil (20 μM, Alomone Labs), a T-type Ca^2+^ channel blocker, from the current recorded before addition of the drug, and the amount of calcium entered was quantified by integrating 100 ms calcium current from the onset of stimulus voltage.

All neurons with an access resistance larger than 25 MΩ after formation of the whole-cell, which indicates insufficient breakage of cell membrane, were discarded. Voltage signals were low-pass filtered at 10 kHz and sampled at 50 kHz, and current signals were low-pass filtered at 2 kHz and sampled at 10 kHz. All experiments were performed at a temperature of ∼33 °C with a temperature controller (TC324B; Warner). The liquid junction potential was calculated to be 15.7 mV for the solution used in this experiment. Data presented in the article were not adjusted by this value. After recording of each neuron, the slice was captured through a 5× objective with the infrared CCD camera, the tip of the electrode was documented and was taken as the position of the recorded neurons.

### Analysis of electrophysiological parameters

All electrophysiological data were analyzed with ClampFit 10.2 (Molecular Devices), MATLAB R2009a (Math Work) and Mini analysis 6.0 (Synaposoft). In total, 29 electrophysiological parameters (underlined) were measured, similar to previous studies (Karagiannis et al., 2009, Fujita et al., 2017). The resting membrane potential (*<u>V</u>*<u>_m_</u>) was measured immediately after the formation of the whole-cell, without current injection. Input resistance (*<u>R</u>*<u>_m_</u>) and the membrane time constant (*<u>τ</u>*<u>_m_</u>) were obtained by averaging the voltage response to a hyperpolarization current step (20 pA, 500 ms, ∼5 repeats). Specifically, *R*_m_ was determined by the voltage response at the steady state to the current while the *τ*_m_ was determined by fitting this averaged voltage response to a single exponential curve. The membrane capacitance (*<u>C</u>*<u>_m_</u>) was calculated according to the formula *C*_m_=*τ*_m_/*R*_m_. Some neurons displayed a “sag” following the initial hyperpolarization during injection of hyperpolarizing current pulses. This is mainly due to hyperpolarization activated cationic current (*I*_h_). To measure *I*_h_, a sag index (<u>Δ*G*_Sag_</u>) was calculated. Specifically, hyperpolarization current pulses from 0 to −100 pA were delivered to the neurons and the inactive sag conductance (*G*_hyp_) was measured as the slope of the linear portion of an *I*–*V* plot, in which *V* was the maximal negative potential at the beginning (0-100 ms) of the hyperpolarizing pulses. The active sag conductance (*G*_sag_) was measured as the slope of the linear portion of an *I*–*V* plot, in which *V* was the average negative potential at the end (700-800 ms) of the hyperpolarizing pulses. Δ*G*_Sag_, quantified as the change in membrane conductance according to 100*(*G*_hyp_ - *G*_sag_)/*G*_sag_, is known to indicate *I*_h._ Occurrence of post-inhibitory rebound <u>(% rebound</u>) were determined by the appearance of depolarization (>3 standard deviation above the baseline) following injection of hyperpolarizing current (100 pA). Rebound amplitude and duration were measured unless the neurons fired action potential (AP) on the rebound. <u>Rebound amplitude</u> was defined as the difference between the rebound peak and the baseline afterwards. <u>Rebound duration</u> was defined as the time interval between the offset of the hyperpolarization to the point of rebound reaching back to the baseline. Occurrence of neurons that fire rebound AP (<u>%rebound AP</u>) were also tallied.

All properties of action potential (AP) were determined by analyzing only the first spike elicited by injection of depolarization current step. <u>Rheobase</u> was the minimum current that elicited spike when holding to −60 mV. <u>Spike latency</u> was computed as the time from the onset of a current pulse to the peak of the AP. <u>Spike d*V*/d*t*_max_</u> was calculated as the maximum derivative of the voltage (d*V*/d*t*) during the first AP. Spike threshold (*<u>V_th_</u>*) was defined as the voltage when d*V*/d*t* equals to 20 V/s. <u>Spike peak</u> was the maximum voltage during the AP, and <u>spike amplitude</u> was measured by subtracting the *V_th_* from the spike peak. <u>Spike half-height duration</u> was defined as the width at half spike amplitude above the threshold. <u>Spike rise time</u> and <u>decay time</u> were calculated as the time between 10-90% of the spike amplitude on either the rise or decay phase of the AP. Occurrence of after-hyperpolarization <u>(% AHP</u>) was defined as the appearance of a trough after the AP down slope with a differentiation > 0 when back to baseline. The <u>AHP amplitude</u> and <u>latency</u> were measured as the difference in amplitude and time between *V_th_* and the most hyperpolarized point of the AHP trough. The <u>AHP duration</u> was measured as the time between the *V_th_* value point during the offset of the AP to the next *V_th_* value point after the AHP or to the plateau of the later rehabilitation.

To describe the variability of neuronal firing patterns around the threshold, the instantaneous firing frequency in response to the minimal current injection that elicited more than three action potentials were plotted and fitted to a linear curve according to the equation *F*_threshold_ = *m*_threshold_ × *t* + *F*_min_, in which *<u>m</u>*<u>_threshold_</u> is the slope termed adaptation, *t* is the time, and *<u>F</u>*<u>_min_</u> is the minimal steady-state frequency. Indeed, we observed variable firing behaviors around threshold with neurons exhibiting bursting firing, or tonic firing, or firing of adapting or irregular trains of action potentials. The percentage of bursting cells (<u>% bursting</u>) were also calculated. Most bursting neurons, including initial bursting and sustained bursting, showed a relatively bigger *m*_threshold_ (*m*_threshold_∣ > 20). The initial bursting neurons exhibit some high-frequency spikes at the beginning the trace, then fire at low frequency in the steady-state, while the sustained bursting neurons exhibit several bunches of high-frequency spikes. Among the non-bursting neurons, we also observed tonic firing (mostly ∣ *m*_threshold_∣ < 4), adapting firing, late spike firing and irregular firing. At higher stimulation intensities, the maximal firing rate (*<u>F</u>*<u>_max_</u>) was measured on the last trace before prominent reduction of the AP amplitude. The <u>spike amplitude accommodation</u> was analyzed on the train with most APs and was measured as the difference between the peak voltage of the AP of the biggest amplitude and the peak voltage of the following AP of the smallest amplitude. In the trace that was used to determine the neuronal firing patterns, amplitudes and half-height duration of the first AP and second AP were measured. Amplitude reduction (<u>Var A</u>) and duration increase (<u>Var D</u>) were quantified as the percentage of reduction and increase normalized to the value of the first AP, respectively.

### Analysis of soma morphological parameters

Soma was manually delineated using Image-Pro Plus 6.0 software (Media Cybernetics) from infrared contrast somatic pictures taken before whole-cell recording. Parameters including soma surface area, length of major and minor axis, perimeter and roundness were extracted.

### Single-cell RT-PCR

Single-cell RT-PCR were performed as previously described (Pfeffer et al., 2013, Li et al., 2019). Briefly, immediately after whole-cell formation, the contents of the neurons were aspirated into the patch electrode. The electrode tip was then broken off to release the content into a thin-wall PCR reaction tube for RT-PCR. cDNA library of the recorded cells was generated with random hexamers using the Superscript IV kit (Invitrogen, #18091050) according to the manufacturer’s protocol. Internal solution was transcribed along as a negative control. Afterwards, multiplex PCR was carried out with 2×Taq PCR Master-Mix (Tiangen, #KT201-02) using primers for targeted genes in a volume of 25 μl. Multiplex primers were designed to amplify exonic DNA sequences and spanned at least one exon-intron boundary. Water control experiments were regularly performed to detect contamination. Next, nested single gene PCR was carried out in a volume of 25 μl with a 1:25 dilution of the multiplex PCR products using the standard 2×Taq PCR Master-Mix (Tiangen, #KT201-02). Nested primers were designed to amplify 100–400 bp DNA sequences within the boundary of the multiplex PCR primers. PCR products were visualized using standard agarose gel electrophoresis and visualized/documented under the UV light. All primers used were first tested with dilutions of mPOA cDNA libraries for sensitivity and specificity. The following primers are used in this study: *Vgat*, multiplex primer (508 bp), 5’-TCCTGAAATCGGAAGGCGAG-3’, 5’-TTGTACATGAGGTTGCCGCT-3’, nested primer (137 bp), 5’-GTCACGACAAACCCAAGATCAC-3’, 5’-GGCGAAGATGATGAGGAACAAC-3’; *Vglut2*, multiplex primer (327 bp), 5’-AAGCTTTGGCATGGTCTGGT-3’, 5’-ATGACAAGGTGAGGGACTGC-3’, nested primer (136 bp), 5’-ATCTGCTAGGTGCAATGG-3’, 5’-TAAGCTGGCTGACTGATG-3’.

### Fluorescent *in situ* hybridization

Animals were anesthetized with 1% pentobarbital sodium (50 mg/kg, i.p.; Merck) and transcardially perfused with DEPC-treated PBS (D-PBS) followed by ice-cold 4% PFA in PBS. Brains were dissected and post-fixed in PFA over night at 4 °C and dehydrated with 30% sucrose in D-PBS. Afterward, brains were sectioned at 20 μm thickness and mounted onto SuperFrost Plus® Slides (Fisher Scientific, Cat. No. 12-550-15). After drying in the air, slides were stored in −80 °C before being processed according to the RNAscope® Multiplex Fluorescent Reagent Kit v2 User Manual (ACD Bio.). Probes against *Vglut2* and *tdTomato* mRNA were ordered from ACD Bio. and used in the experiment. Images were captured under a 40× objective using a confocal microscope (C2; Nikon).

### Cannula Implantation

Adult mice were anesthetized using isoflurane (0.8–5%) and placed onto a stereotactic frame (David Kopf Instrument, Model 1900). The skull was exposed with a small incision and holes were drilled in the skull to implant cannulas. The coordinate of bregma: AP: +0.000 mm, ML: ±0.450 mm, DV: −4.100 mm was used to target the mPOA bilaterally. Bilateral cannulas (O.D. 0.41 mm × I. D. 0.25 mm, length 4.5 mm, interval 1.0 mm; RWD Inc.) were inserted through the drilled holes and secured onto the skull with dental cement (Super-Bond C&B). Cannula implanted animals were allowed at least one week to recover before behavioral tests.

### Behavioral assays

Prior to the cannula implantation, adult male animals (∼2 months old) were single-housed and were paired overnight with a hormonal primed ovariectomized C57BL/6 female twice. Mating behavior tests were carried out >1 week after the cannula implantation surgery. All behavioral tests were initiated at least 4 hours after the onset of the dark cycle. Briefly, 20 min before the mating behavioral tests, 500 nl of saline or mibefradil (1 mg/ml in saline, Alomone Labs) was infused per side at a rate of 0.15 μl/min through a fitted stainless steel injector that protrudes 1 mm beyond the tip of the cannula (RWD). The injector was left in place for 1 min after infusion. Afterwards, a hormonal primed ovariectomized C57BL/6 female was introduced to the cage and behaviors were videotaped for ∼30 min using an infrared camera at a frame rate of 25 Hz. Each animal was tested once a week and 2-3 trials each for drug or saline infusion on a balanced delivery schedule.

When the last behavioral trial was finished, a fluorescent dye Fluor594 (200 μM, Invitrogen) was delivered through the cannulas. Animals were then anesthetized with 1% pentobarbital sodium (50 mg/kg, i.p.; Merck), brains were dissected and post-fixed in 4% paraformaldehyde (PFA) in PBS over night at 4 °C. Afterwards, brains were sectioned at 40 μm thickness using a vibratome (VT1000S, Leica). Brain sections were stained DAPI (5 mg/ml, 1:1000, Sigma) in AT (0.1% Triton and 2 mM MgCl2 in PBS) for 10 min at room temperature. After several washes in PBS, brains sections were mounted onto glass slides. Images were captured under a 10× objective using a fluorescent microscope (VS120; Olympus) and analyzed to verify correct targeting of the mPOA. Only the animals with bilateral hits of mPOA were processed for video analysis. In total, twenty males that were implanted with cannulas, of which three were excluded for mis-targeting, one died prior to behavioral testing, the cannula in one was clogged, and video system malfunction happened during videotaping of one. Thus, behavioral results of fourteen animals were analyzed.

Videos were manually annotated on a frame-by-frame basis using a custom written MATLAB program as previously described (Xu et al., 2012, Wei et al., 2018), by experimenters blinded to the group information. Behaviors were scored according to the following criteria: nose-to-urogenital contact is defined as “sniff”, also known as “chemoinvestigation”. “Mount” is scored when the animal places its forelimbs on the back of the stimulus, climbs on top and moves the pelvis. Rhythmic pelvic movements after mount are scored as “intromit.”

### Statistical analysis

Data were statistical analyzed and plotted with GraphPad Prism 7. Values are presented as mean ± SEM. In all cases of electrophysiological parameters, *N* refers to the number of neurons. For comparison of two groups, if *N* < 30, the data were first tested to be Gaussian distribution with Shaprio– Wilk normality test, and if the variances were tested to be equal with *F* test, *p* values were calculated with paired or unpaired two-tailed Student’s *t* test; otherwise, unpaired *t* test with Welch’s correction was applied to data with Gaussian distribution but unequal variances. For all other cases (*N* > 30 or *N* < 30 but not normally distributed), data were analyzed using a Mann–Whitney *U* test. Categorical data were compared with Fisher’s exact test. \**p* < 0.05, \*\**p* < 0.01, \*\*\**p* < 0.001.

### Code Availability

Matlab codes used for data analysis are available upon request.

## Results

### Whole-cell patch clamp recording of mPOA neurons

We performed whole-cell patch clamp recordings of mPOA neurons in acute brain slices cut from adult animals (Figure 1A). Before recording, infrared contrast somatic pictures were taken for analysis of soma morphology (Figure 1A). Positive and negative current steps were injected into the recorded neurons to obtain membrane responses (Figure 1B, for details see “Methods”), from which 29 electrophysiological features related to passive membrane properties, rebound, and action potential (AP) were extracted. Afterwards, spontaneous EPSCs and IPSCs were recorded under voltage clamp mode with membrane potential held at −70 mV and 0 mV respectively (Figure 1C). At the end of the recording, we took pictures of the recording electrode, from which the coordinates of the patched neuron, was estimated based on relative positions of the tip of the electrode to anatomical landmarks such as the third ventricle and the commissure (Figure 1A). In total, we sampled 111 neurons from 16 males and 85 neurons from 14 females of C57BL/6 background. Projected positions of these neurons in a reference brain atlas show that they are tightly clustered within the mPOA as defined by the atlas and there is no systematic deviation between the two sexes (Figure 1D).

**Figure 1.**
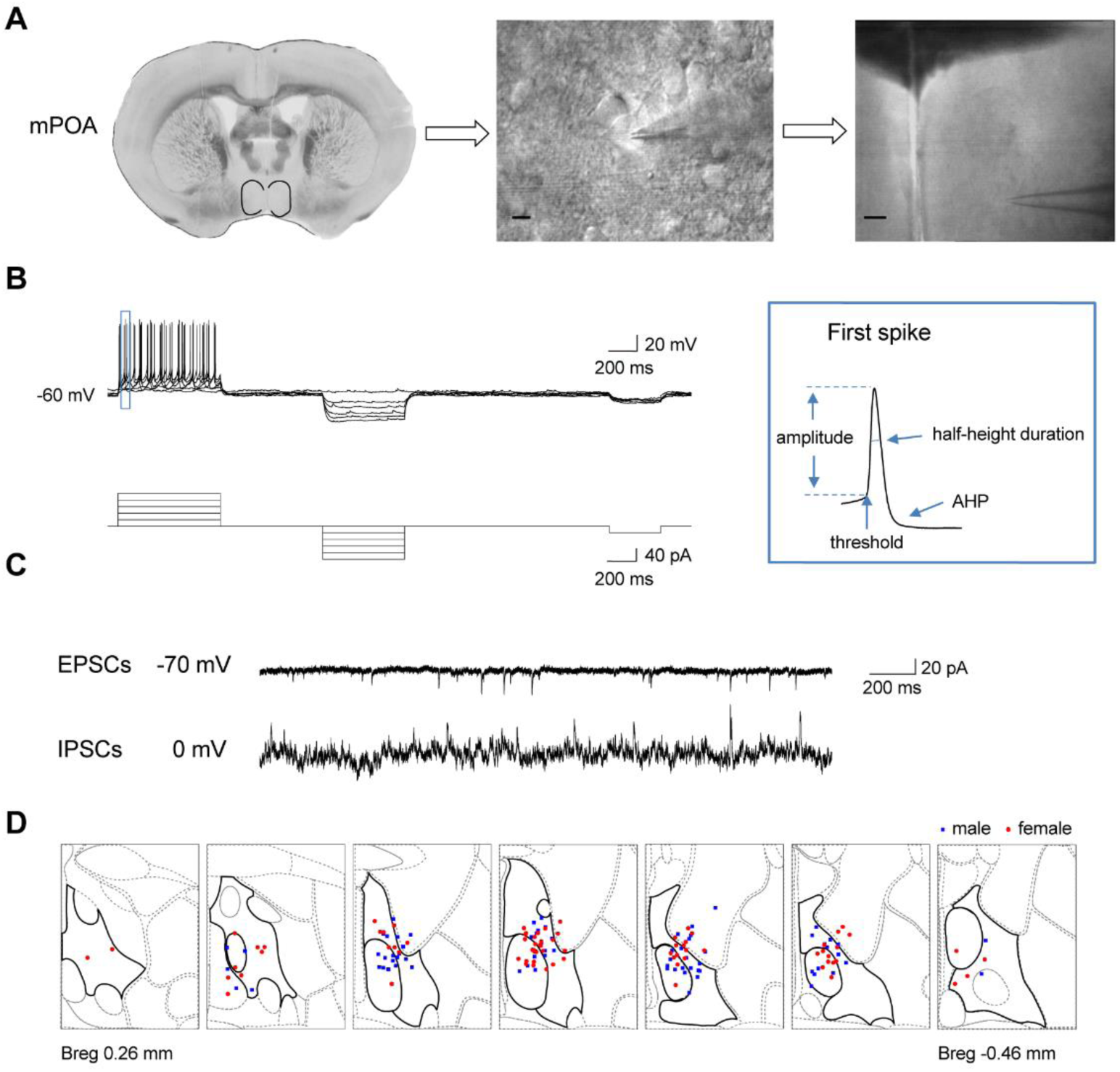
Slice recordings of mPOA neurons. **A.** Acute brain slices containing the mPOA, as marked out by the black lines, were chosen. Pictures of neurons were captured under a 40× objective just before the electrode reached the target neuron (scale bar, 10 μm) and were used to analyze the soma morphology parameters of the recorded neurons. After recordings, the tip of the electrode was visualized under a 5× objective (scale bar, 100 μm) to document the coordinates of the recorded neuron. **B.** Example voltage signals (top traces) recorded from a mPOA neuron in response to stimulation currents (bottom traces). The first spike elicited in this neuron as highlighted in the blue square on the right was processed to calculate electrophysiological parameters related to the action potential, including threshold, amplitude, half-height duration and after-hyperpolarization (AHP). **C.** Representative traces of spontaneous excitatory postsynaptic currents (EPSCs; top) and inhibitory postsynaptic currents (IPSCs; bottom). **D.** Positions of all neurons recorded in the wildtype animals were mapped onto the reference mouse brain atlas from the Allen Brain Institute (from bregma 0.26 mm to −0.46 mm). The mPOA was highlighted with thick black lines in each section.

### Significant sex differences in electrophysiological properties of mPOA neurons

By comparing mPOA neurons recorded in wildtype male and female mice, we found that male mPOA neurons had larger soma surface area and longer minor axis than female, which indicates bigger mPOA neurons (Table 1). Similarly, we observed significant sex differences in passive membrane properties (Table 2), with the resting membrane potential (*V*_m_) being more positive on average in male than female (male, −55 ± 0.62 mV, female, −57.28 ± 0.78 mV, Mann–Whitney *U* test, *p*=0.0293, *U*=3862), and the input resistance (*R*_m_) and membrane time constant (*τ*_m_) being smaller in female than male (*R*_m_, male, 509.67 ± 21.43 MΩ, female, 658.35 ± 29.72 MΩ, Mann–Whitney *U* test*, p*<0.0001, *U*=3229; *τ*_m_, male, 26.16 ± 1.15 ms, female, 30.19 ± 1.42 ms, Mann–Whitney *U* test*, p*=0.0302, *U*=3865). By contrast, electrophysiological properties that relate to action potential were comparable between male and female.

**Table 1.** Soma properties of mPOA neurons in wildtype male and female mice.

|  | male<br>(N=102) | female<br>(N=83) | *p* value | Mann-Whitney<br>*U* |
| --- | --- | --- | --- | --- |
| (1) soma surface<br>area (μm<sup>2</sup>) | 178.02 ± 3.84 | 164.29 ± 4.1 | 0.0061 | 3243 |
|  | female << male. |  |  |  |
| (2) major axis (μm) | 19.50 ± 0.32 | 19.02 ± 0.35 | 0.1254 | 3677 |
| (3) minor axis (μm) | 12.15 ± 0.21 | 11.52 ± 0.22 | 0.0427 | 3499 |
|  | female < male. |  |  |  |
| (4) perimeter (μm) | 56.05 ± 0.86 | 54.55 ± 1.02 | 0.0873 | 3613 |
| (5) roundness | 1.42 ± 0.02 | 1.46 ± 0.03 | 0.3817 | 3915 |

5 All variables are described by the mean ± SEM. *N*, Number of cells. < indicates 6 significantly smaller with *p* < 0.05; << indicates significantly smaller with *p* < 0.01. Mann– 7 Whitney *U* test was performed between wildtype male and female mice.

**Table 2.**
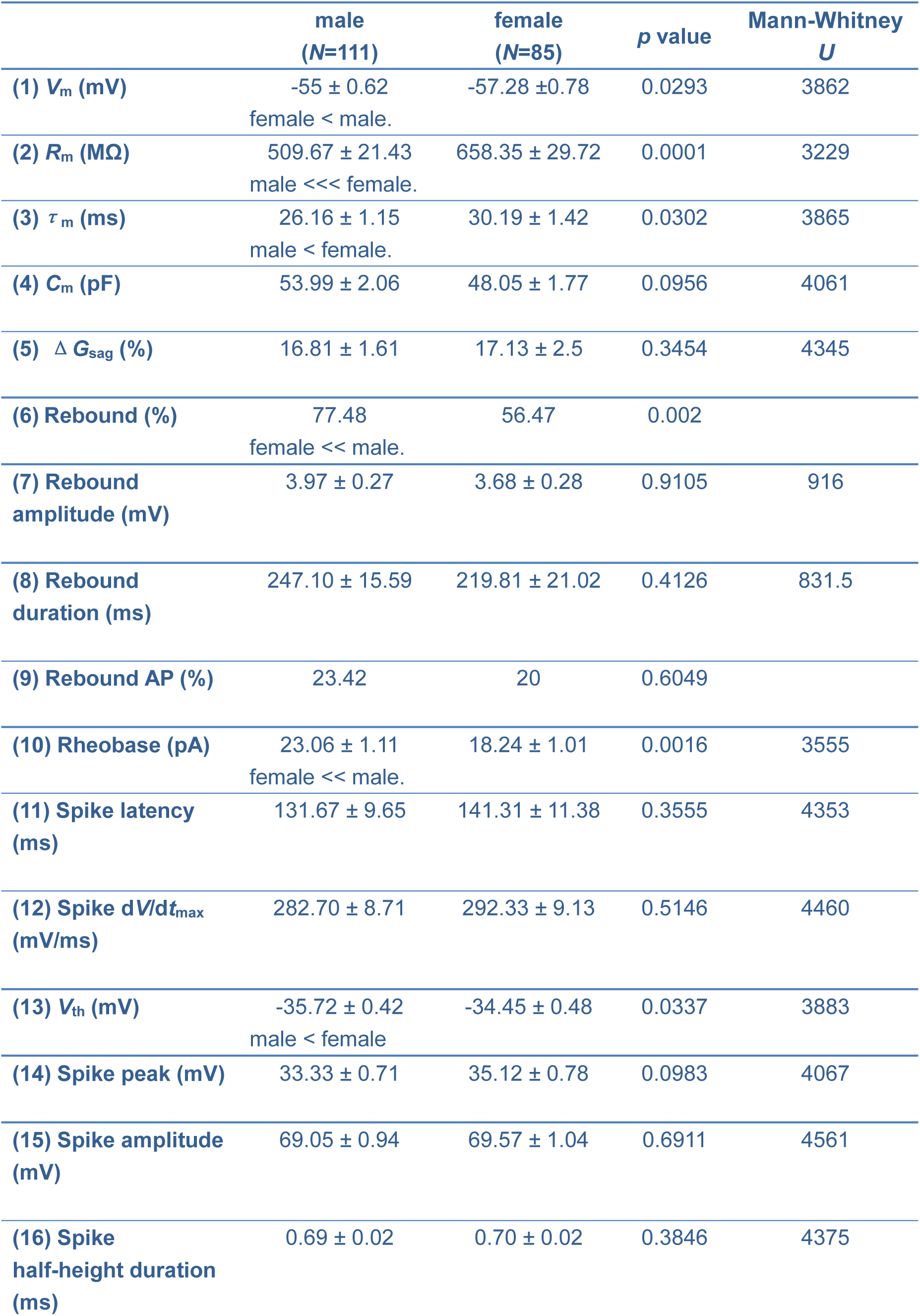
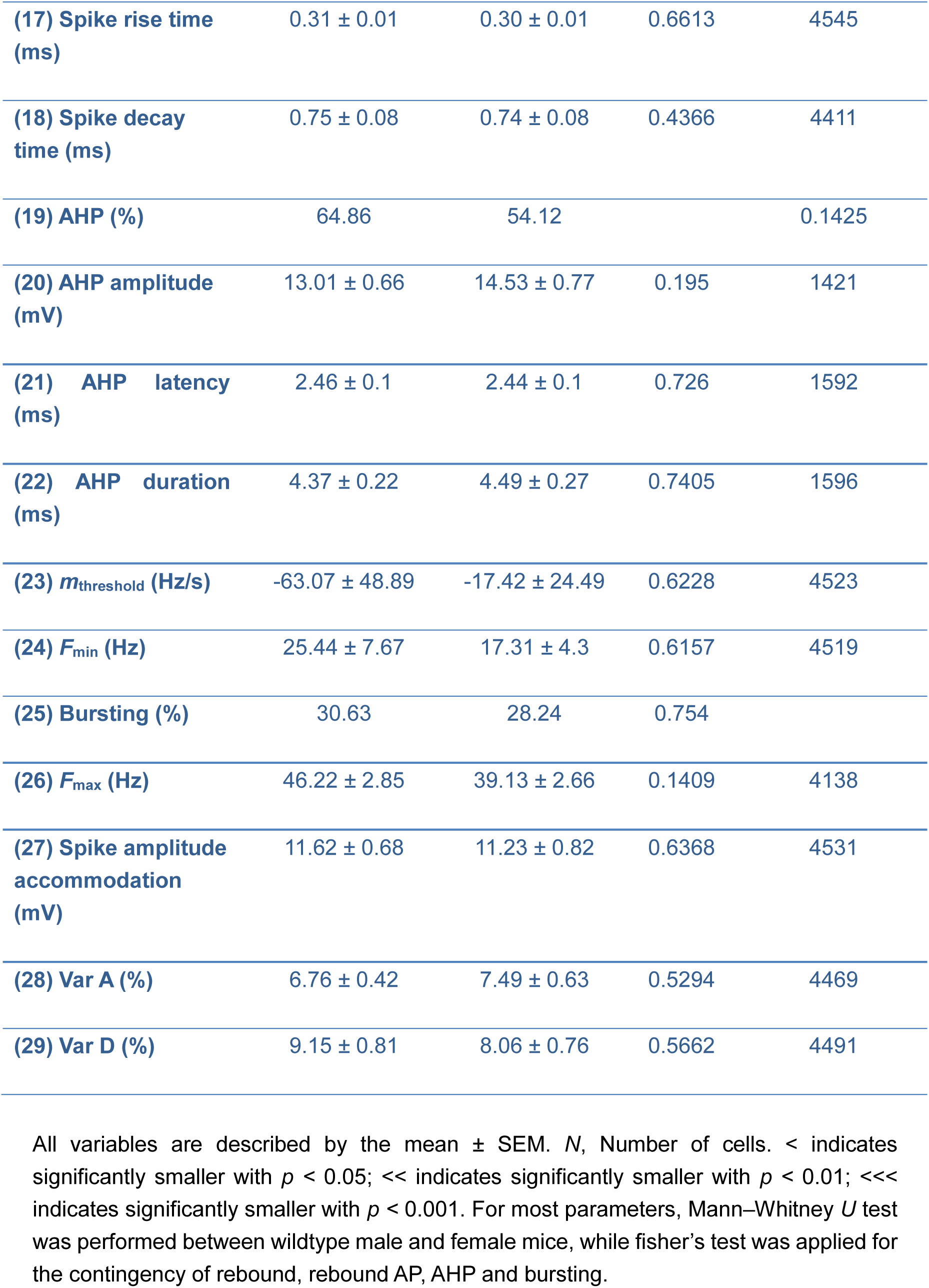
Electrophysiological properties of mPOA neurons in wildtype male and female mice.

Interestingly, even though the minimum current injection needed to produce an action potential, known as rheobase, appeared to be larger in male mPOA neurons than female (male, 23.06 ± 1.11 pA, female, 18.24 ± 1.01 pA, Mann–Whitney *U* test*, p*=0.0016, *U*=3555), the voltage threshold for an action potential (*V_th_*) was actually lower in males (male, −35.72 ± 0.42 mV, female, −34.45 ± 0.48 mV, Mann–Whitney *U* test*, p*=0.0337, *U*=3883). Considering that the measured rheobase was acquired by holding the membrane potential at −60 mV, we therefore calculated the theoretical rheobase, defined as (*V*_th_-*V*_m_)/*R*_m,_ and found no sex difference in this value (male: 45.03 ± 2.32 pA, female: 40.24 ± 2.19 pA; Mann–Whitney *U* test, *p* = 0.1184, *U* = 4064), suggesting that male and female mPOA neurons require similar depolarization current to be excited. On another hand, the percentage of neurons that rebound after injection of hyperpolarization current was significantly higher in male than female (male, 86 out of 111, female, 48 out of 85, Fisher’s exact test, *p*=0.002), implying higher neuronal excitability in males under certain conditions.

Zooming out from single action potential, we observed variable firing patterns around the threshold that can be broadly divided into non-bursting and bursting with the later includes both initial bursting and sustained bursting (Figure 2A-B). These firing patterns were observed with similar proportion in male and female (Figure 2C) with the predominant type being non-bursting cells (male, 69.37%; female,71.76%). As expected, bursting and non-bursting cells differ significantly in AP decay time, AHP amplitude and duration, *m*_threshold,_ *F*_min_, Var A and Var D, with bursting cells in male also displaying wider AP half-height duration, lower *F*_max_ and larger AP amplitude accommodation than non-bursting cells, and bursting cells in female displaying more depolarized *V_m_* and smaller spike d*V*/d*t*_max_ (Table 3, Supplementary Table 3-1). More importantly, when we compared electrophysiological properties of bursting neurons between male and female, we essentially found no sex differences except for *V_th_* (Table 3, Supplementary Table 3-1). By contrast, nearly all identified sex differences were found in non-bursting cells (Table 3, Supplementary Table 3-1), consistent with a higher proportion of these cells in the mPOA (Figure 2C).

**Figure 2.**
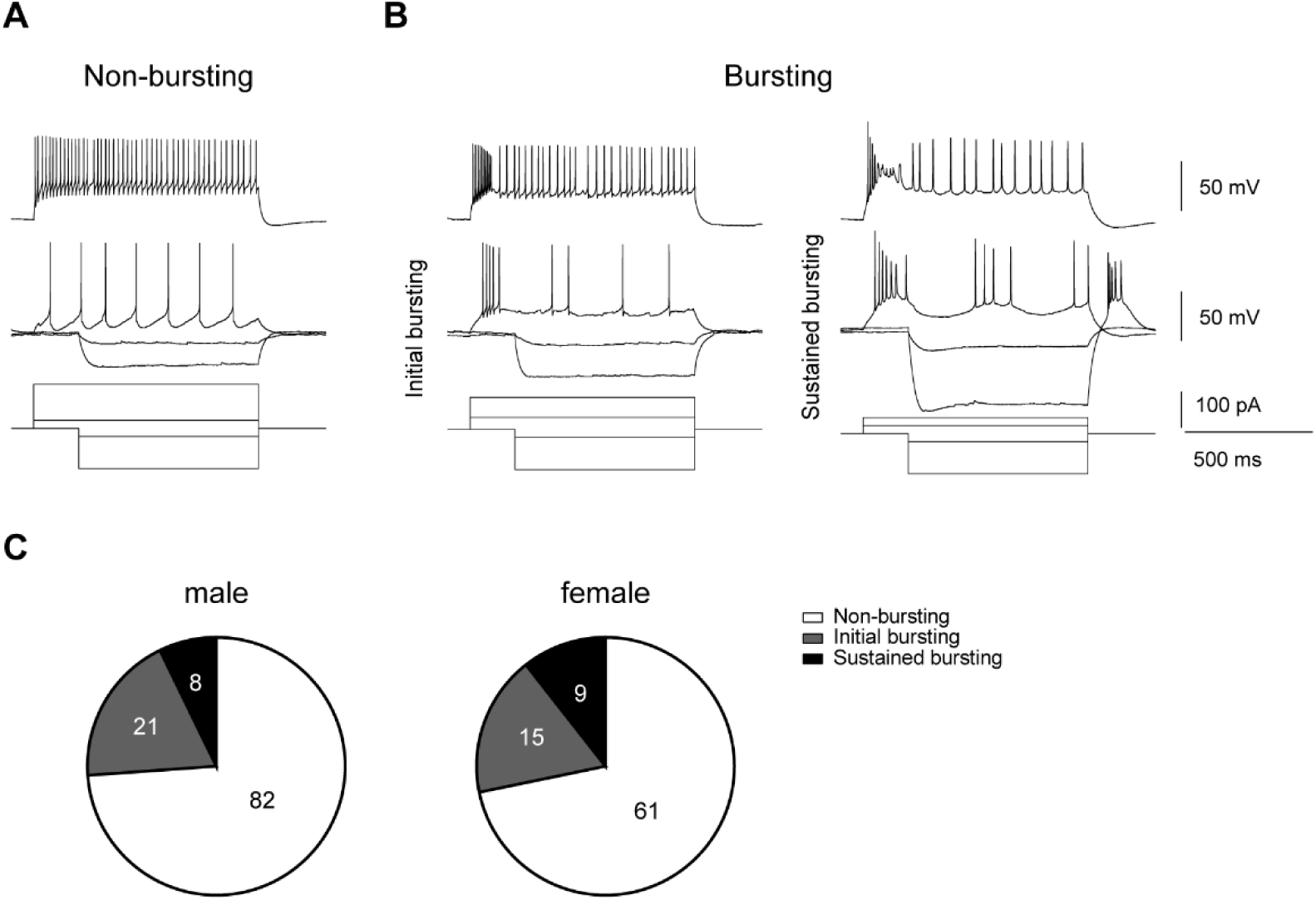
Firing patterns of mPOA neurons. **A-B**. Representative traces of a non-bursting (A) and two bursting neurons (B, initial bursting on the left and sustained bursting on the right). Top traces show neuronal responses to depolarizing current step injections (1000 ms) that could elicit saturated firing of action potentials. Middle traces show neuronal responses to the minimal around threshold depolarizing current step injections (1000 ms) that could elicit more than three APs along with neuronal responses to two hyperpolarizing current step injections (800ms). Bottom traces show the stimulus currents used for the corresponding neuron. **C.** The percentage of initial bursting, sustained bursting and non-bursting mPOA neurons in male and female.

**Table 3.**
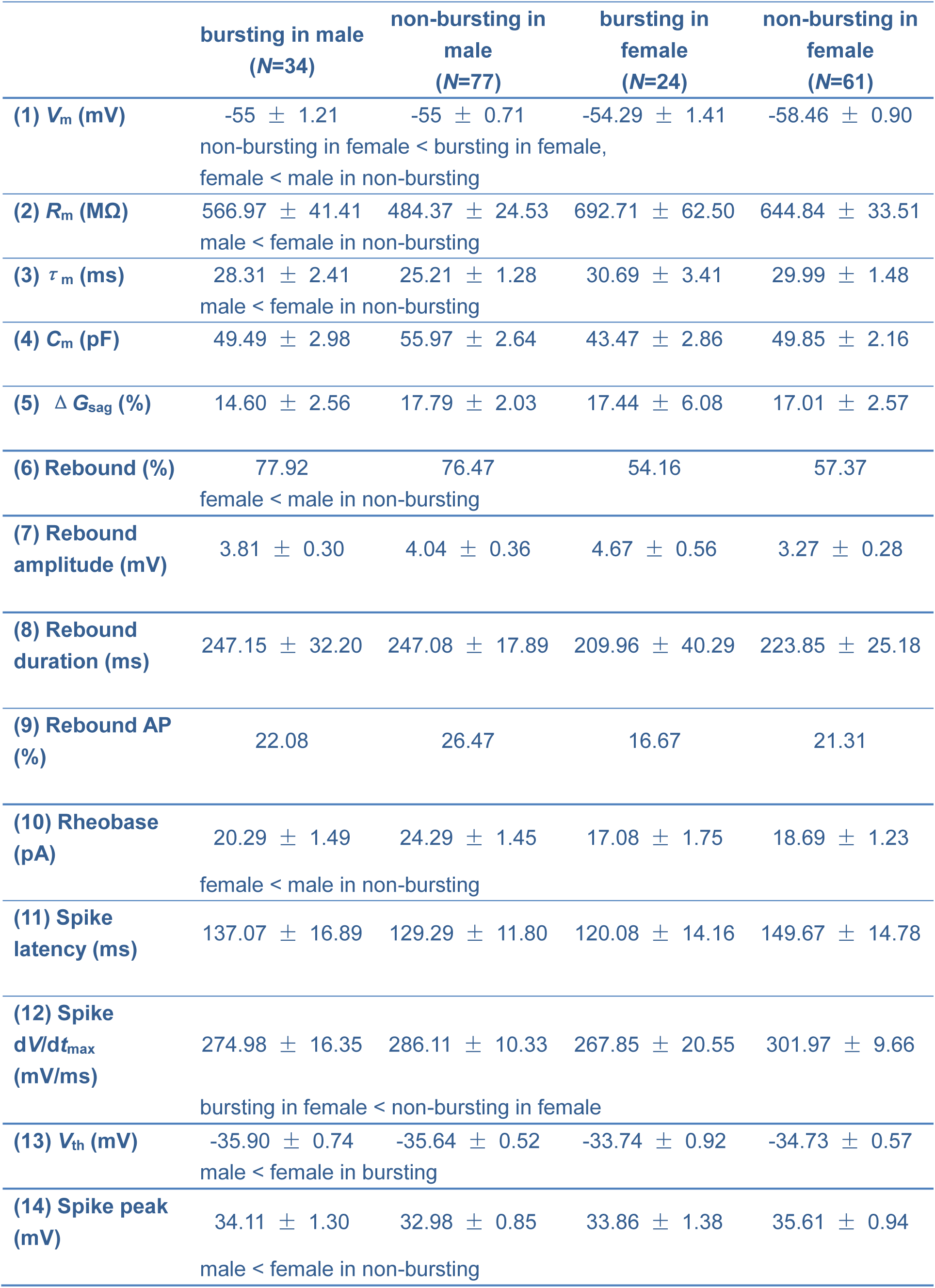
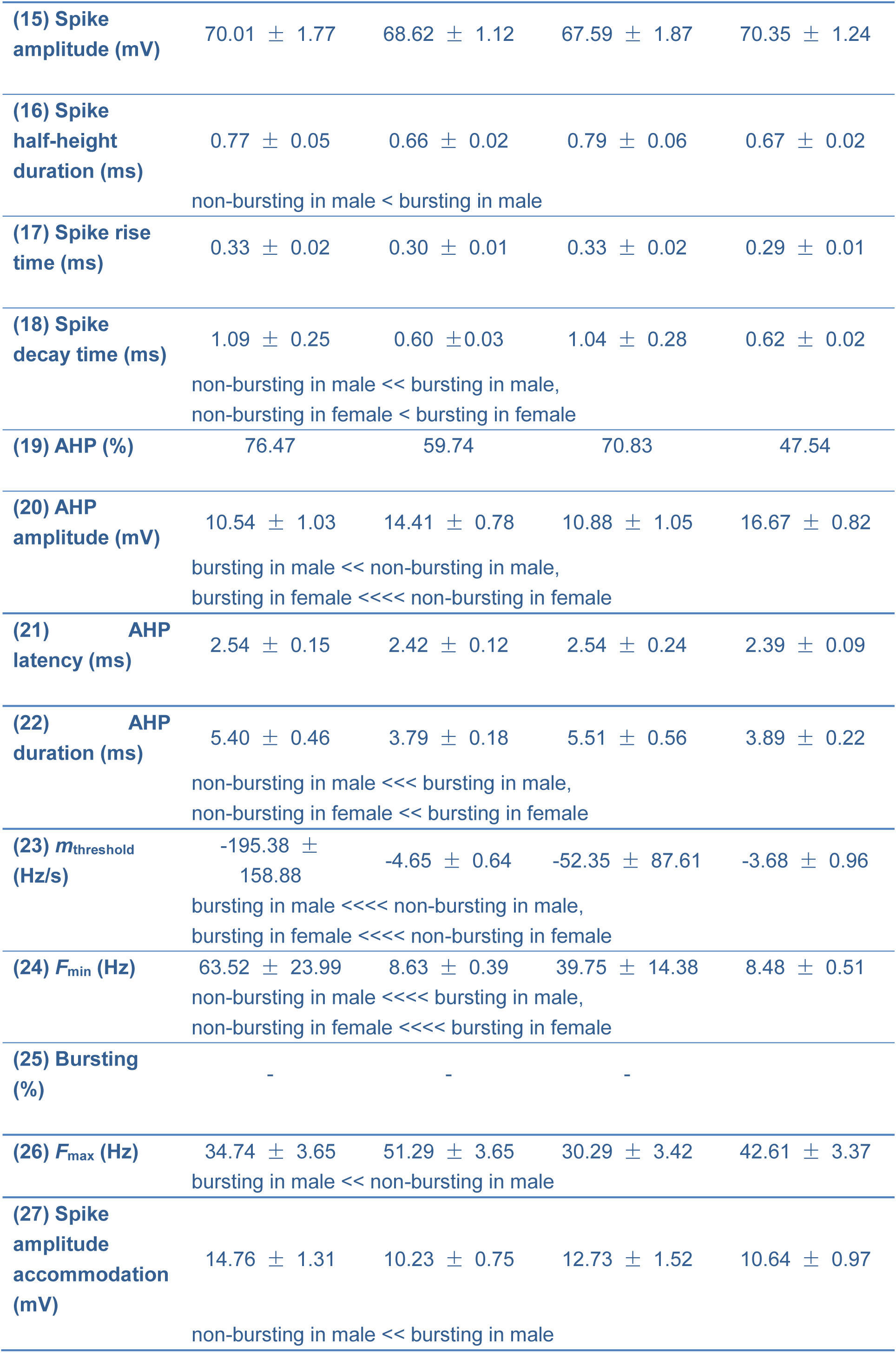
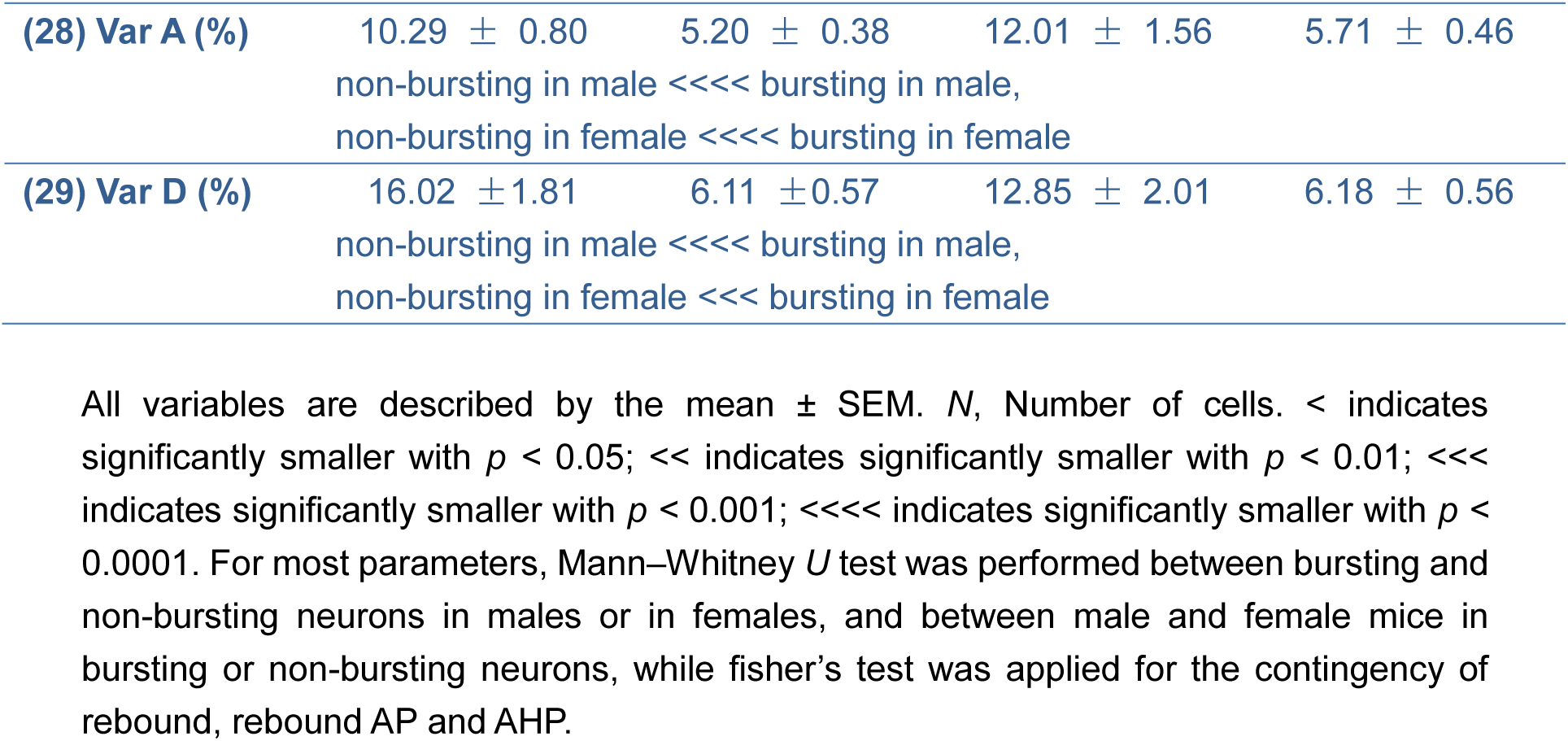
Electrophysiological properties of bursting and non-bursting neurons in wildtype male and female mice.

### No sex differences in spontaneous postsynaptic currents of mPOA neurons

After the current clamp recording of membrane properties, we recorded spontaneous excitatory and inhibitory postsynaptic currents (sEPSCs and sIPSCs) of mPOA neurons. In total, 89 and 60 out of 111 mPOA neurons patched in male were measured for sEPSCs and sIPSCs respectively, and 65 and 54 out of 85 mPOA neurons in female were measured for sEPSCs and sIPSCs respectively. Representative traces of sEPSCs and sIPSCs are shown in Figure 1C. No significant sex differences were observed in either frequency or amplitude of sEPSCs or sIPSCs (Table 4), suggesting that male and female mPOA neurons receives similar amount of total excitatory and inhibitory inputs.

**Table 4.**
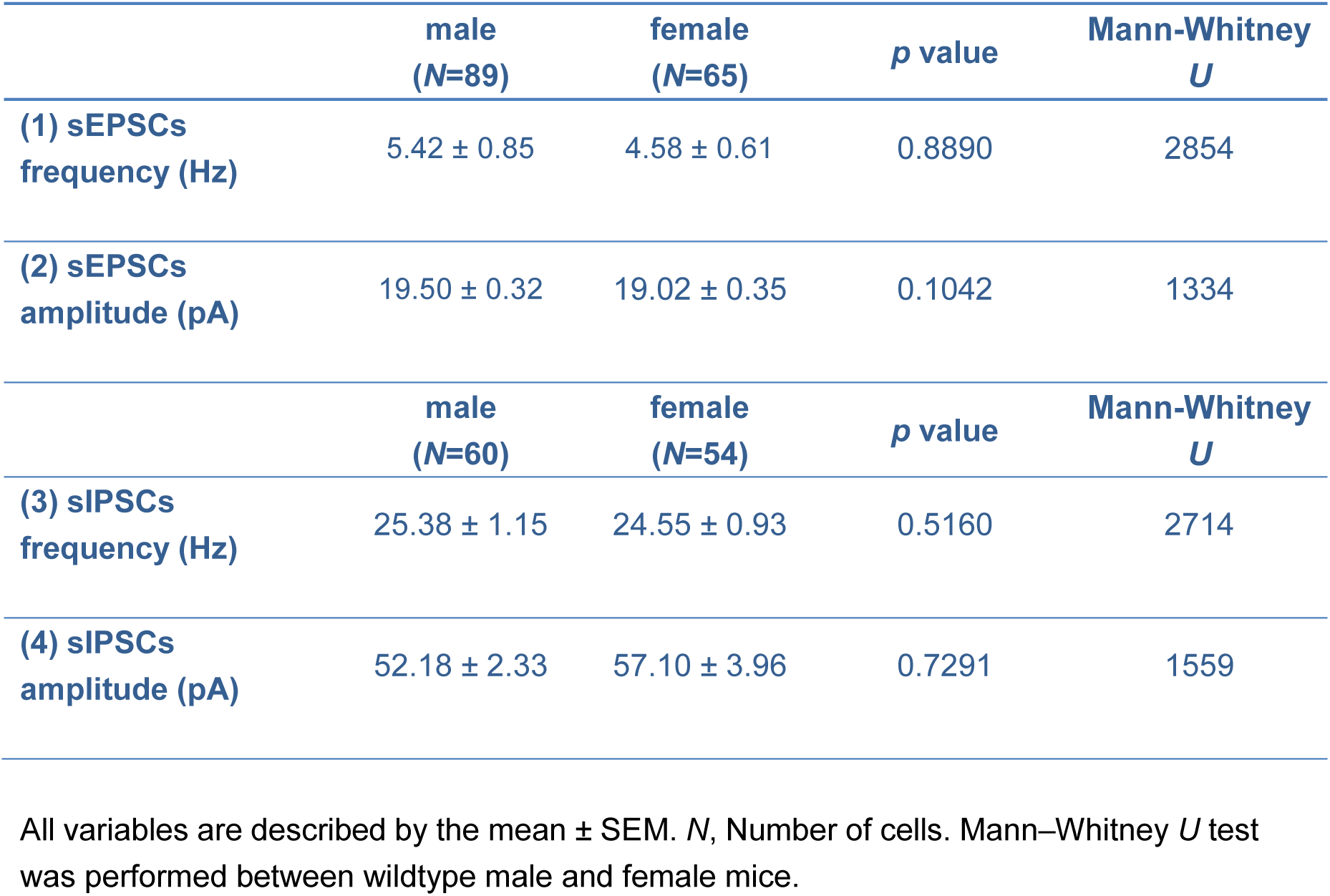
Spontaneous excitatory and inhibitory inputs to mPOA neurons in wildtype male and female mice.

### Sex differences in electrophysiological properties depend on cell type

Single-cell RT-PCR revealed that ∼80.95% of mPOA neurons surveyed unbiasedly with patch-clamp expressed *Vgat* (vesicular GABA transporter), a fraction of which also co-expressed *Vglut2* (Figure 3A), consistent with existing results from *in situ* experiments and from single-cell transcriptome analysis of this region (Moffitt et al., 2018, Wei et al., 2018). Thus, to more selectively investigate electrophysiological properties of mPOA *Vglut2+* neurons, we crossed *Vglut2-Ires-Cre* with *Ai9* to create a compound mouse line (*Vglut2::Ai9*) that fluorescently labels Vglut2+ neurons, as confirmed by >95% co-localization of *tdTomato* and *Vglut2* in this line (Figure 3B). Using this mouse line, we recorded 56 *tdTomato*+ neurons in the mPOA of 10 male mice and 63 *tdTomato*+ neurons in 9 female mice (Figure 3C).

**Figure 3.**
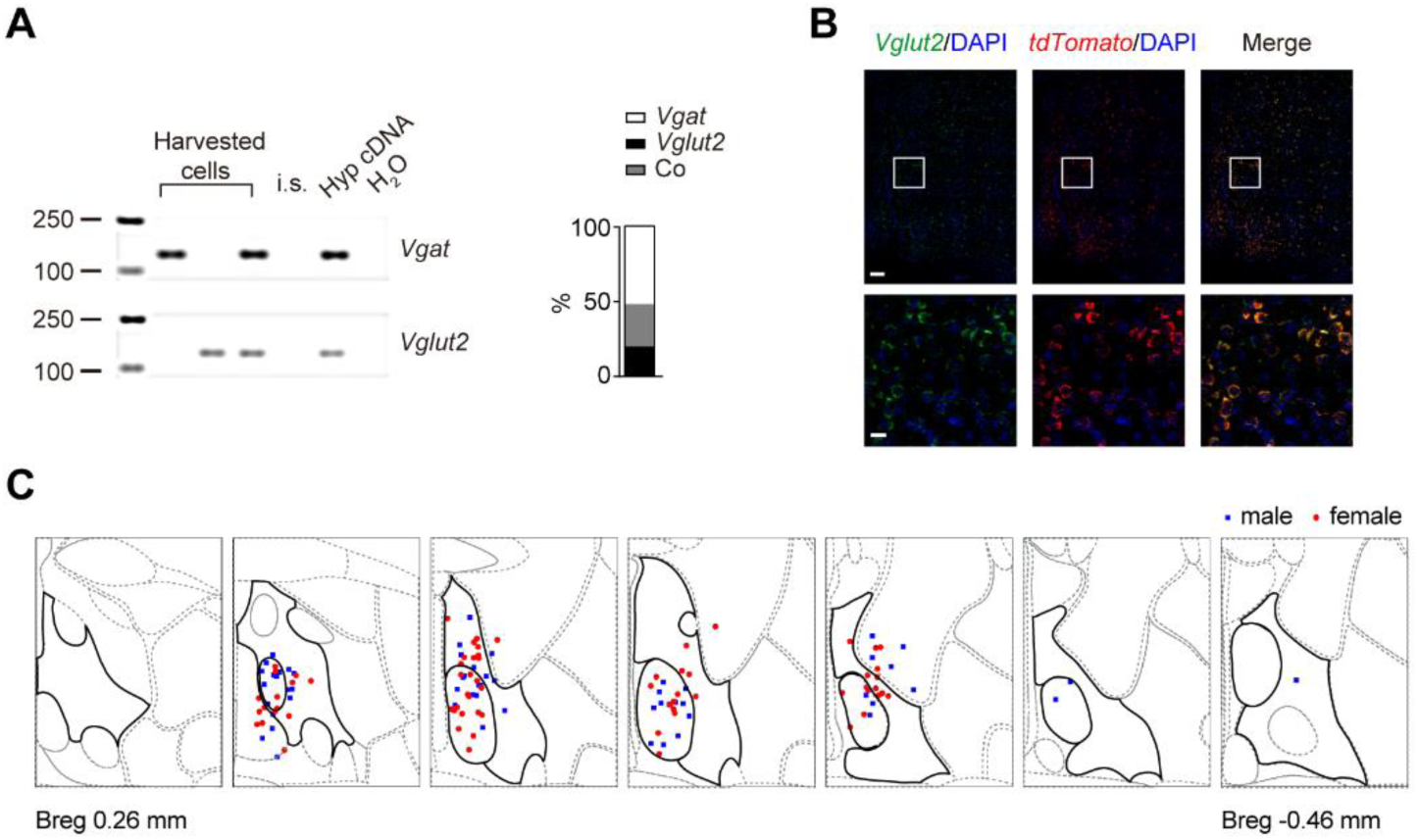
Slice recording of mPOA *Vglut2*+ neurons. **A**. Constitution of *Vgat+* and *Vglut2+* neurons in unbiased patch of the mPOA. Left panel: a representative gel image showing single-cell RT-PCR analysis of mPOA neurons with the genes indicated on the right. H_2_O and internal solution (i.s.) served as negative controls and diluted hypothalamus cDNA (Hyp cDNA) as the positive control. Maker size are indicated on the left. Right panel: the percentage of *Vgat+*, *Vglut2+* neurons and neurons that co-expressed *Vgat+* and *Vglut2+* (Co). In total, out of 33 neurons patched, 21 were successfully reverse transcribed, of which 11 were *Vgat*+, 4 were *Vglut2*+ and 6 for both. **B.** Example images showing co-localization of *tdTomato* and *Vglut2* signals in *Vglut2::Ai9* mouse. Images on the bottom are blowups of the corresponding white squares in the images on the top. Scale bar, 100 μm, top; 20 μm bottom. 100% of *tdTomato+* cells are *Vglut2+* and account for ∼95.4% of total *Vglut2+* cells. *N*=3 animals (2 males and 1 female). **C.** Positions of all neurons recorded in *Vglut2::Ai9* animals were mapped onto the reference mouse brain atlas from the Allen Brain Institute (from bregma 0.26 mm to −0.46 mm). The mPOA was highlighted with thick black lines in each section.

Interestingly, all electrophysiological and soma parameters that previously show sex differences (Table 1&2) were now comparable between male and female *Vglut2+* neurons except that male *Vglut2+* neurons were still larger in soma surface area than female (Table 5&6), consistent with the notion that previously observed sex differences mainly derive from *Vgat+* neurons. Instead, male mPOA *Vglut2+* neurons were more likely to burst than female (male, 28 out of 56, female, 19 out of 63, Fisher’s exact test, *p*=0.03825) and showed higher AP amplitude accommodation than female (male, 13.31 ± 0.99 mV, female, 9.23 ± 0.67 mV, *p*=0.0019, *U*=1186). Taken together, these results demonstrate that mPOA *Vgat+* and *Vglut2+* neurons display distinct patterns of sex differences in electrophysiological properties (Figure 5), corroborating the idea that sex differences in the mPOA depend on cell type (Moffitt et al., 2018, Wei et al., 2018).

**Table 5.**
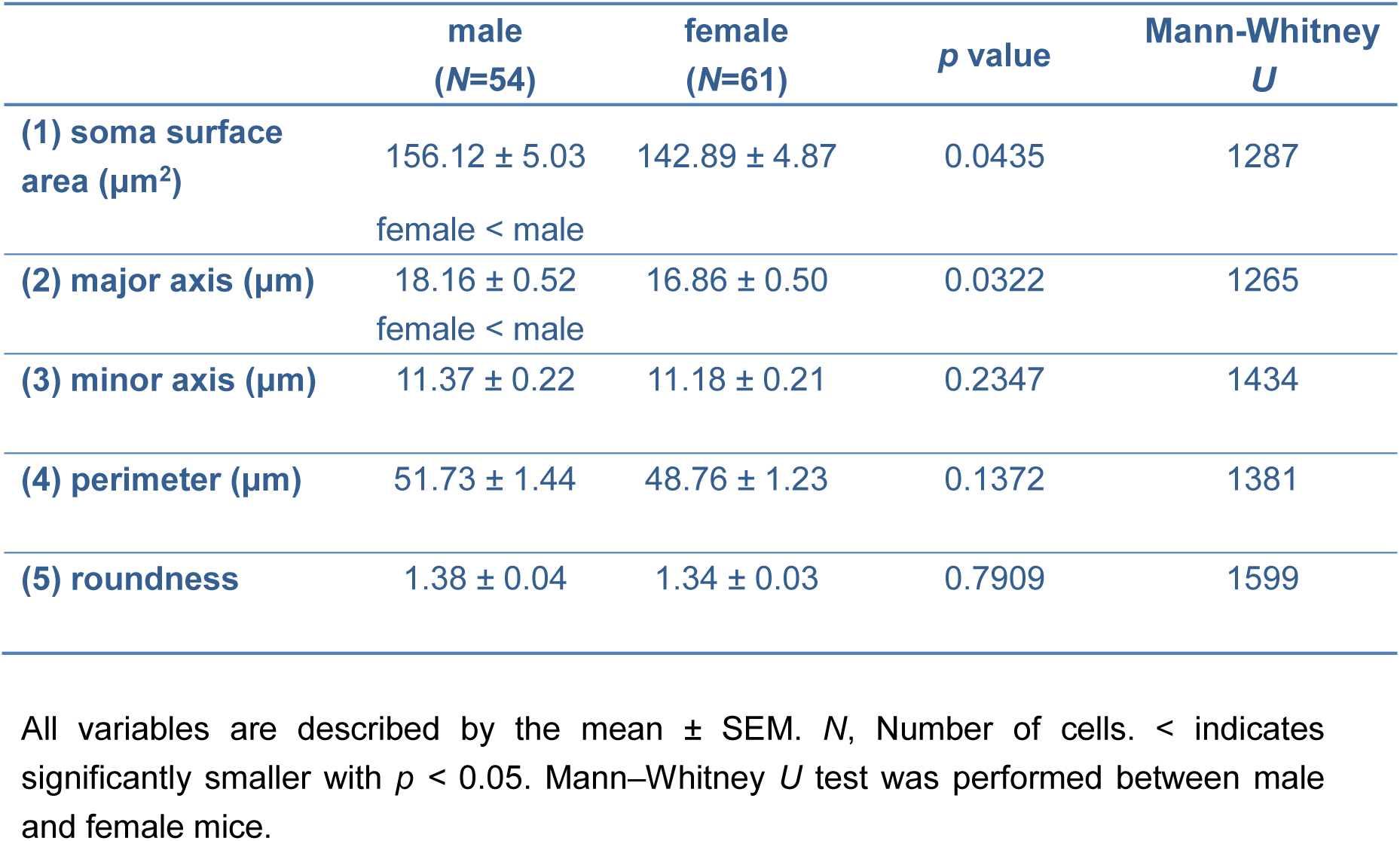
Soma properties of *Vglut2*+ neurons in male and female mice of the *Vglut2::Ai9* line.

|  | male<br>(N=54) | female<br>(N=61) | *p* value | Mann-Whitney<br>*U* |
| --- | --- | --- | --- | --- |
| (1) soma surface<br>area (μm<sup>2</sup>) | 156.12 ± 5.03 | 142.89 ± 4.87 | 0.0435 | 1287 |
|  | female < male |  |  |  |
| (2) major axis (μm) | 18.16 ± 0.52 | 16.86 ± 0.50 | 0.0322 | 1265 |
|  | female < male |  |  |  |
| (3) minor axis (μm) | 11.37 ± 0.22 | 11.18 ± 0.21 | 0.2347 | 1434 |
| (4) perimeter (μm) | 51.73 ± 1.44 | 48.76 ± 1.23 | 0.1372 | 1381 |
| (5) roundness | 1.38 ± 0.04 | 1.34 ± 0.03 | 0.7909 | 1599 |

All variables are described by the mean ± SEM. *N*, Number of cells. < indicates significantly smaller with *p* < 0.05. Mann–Whitney *U* test was performed between male and female mice.

**Table 6.** Electrophysiological properties of *Vglut2*+ neurons in male and female mice of the *Vglut2::Ai9* line.

|  | male<br>(*N*=56) | female<br>(*N*=63) | *p* value | Mann-Whitney<br>*U* |
| --- | --- | --- | --- | --- |
| (1) $V_m$ (mV) | -56.93 ± 1.00 | -57.54 ± 0.97 | 0.6674 | 1683 |
| (2) $R_m$ (MΩ) | 652.82 ± 31.85 | 710.22 ± 37.89 | 0.3597 | 1591 |
| (3) $\tau_m$ (ms) | 29.22 ± 1.52 | 28.53 ± 1.45 | 0.8757 | 1734 |
| (4) $C_m$ (pF) | 46.54 ± 1.85 | 42.02 ± 1.66 | 0.1006 | 1455 |
| (5) $\Delta G_{sag}$ (%) | 9.73 ± 1.57 | 14.90 ± 2.56 | 0.4431 | 1619 |
| (6) Rebound (%) | 42.86 | 52.38 | 0.3591 |  |
| (7) Rebound<br>amplitude (mV) | 4.24 ± 0.70 | 3.27 ± 0.31 | 0.5292 | 171 |
| (8) Rebound<br>duration (ms) | 269.53 ± 34.57 | 210.02 ± 25.32 | 0.1737 | 144 |
| (9) Rebound AP (%) | 16.07 | 11.11 | 0.5915 |  |
| (10) Rheobase (pA) | 18.39 ± 1.24 | 18.57 ± 1.17 | 0.9316 | 1749 |
| (11) Spike latency<br>(ms) | 155.96 ± 20.37 | 143.28 ± 16.41 | 0.5506 | 1651 |
| (12) Spike $dV/dt_{max}$<br>(mV/ms) | 295.88 ± 10.42 | 289.96 ± 10.08 | 0.5866 | 1661 |
| (13) $V_{th}$ (mV) | -33.02 ± 0.77 | -32.94 ± 0.6 | 0.7362 | 1700 |
| (14) Spike peak (mV) | 34.97 ± 0.99 | 35.40 ± 0.82 | 0.7654 | 1708 |
| (15) Spike amplitude<br>(mV) | 67.99 ± 1.17 | 68.33 ± 0.96 | 0.7817 | 1712 |
| (16) Spike<br>half-height duration<br>(ms) | 0.68 ± 0.02 | 0.69 ± 0.02 | 0.9736 | 1758 |
| **(17) Spike rise time (ms)** | 0.30 ± 0.01 | 0.30 ± 0.01 | 0.6225 | 1671 |
| **(18) Spike decay time (ms)** | 0.74 ± 0.10 | 0.62 ± 0.02 | 0.3774 | 1598 |
| **(19) AHP (%)** | 60.71 | 58.73 | 0.8534 |  |
| **(20) AHP amplitude (mV)** | 15.51 ± 1.23 | 15.90 ± 0.86 | 0.9954 | 628 |
| **(21) AHP latency (ms)** | 2.70 ± 0.30 | 2.75 ± 0.13 | 0.0805 | 477 |
| **(22) AHP duration (ms)** | 5.34 ± 0.4 | 4.87 ± 0.31 | 0.3257 | 543 |
| **(23) $m_{\text{threshold}}$ (Hz/s)** | -279.28 ± 177.81 | -75.38 ± 54.12 | 0.8590 | 1730 |
| **(24) $F_{\text{min}}$ (Hz)** | 90.28 ± 58.53 | 24.47 ± 6.48 | 0.6236 | 1671 |
| **(25) Bursting (%)** | 50<br>female < male | 30.16 | 0.0385 |  |
| **(26) $F_{\text{max}}$ (Hz)** | 44.41 ± 4.64 | 46.02 ± 3.16 | 0.2063 | 1526 |
| **(27) Spike amplitude accommodation (mV)** | 13.31 ± 0.99<br>female << male | 9.23 ± 0.67 | 0.0019 | 1186 |
| **(28) Var A (%)** | 7.95 ± 0.58 | 8.26 ± 0.77 | 0.9134 | 1743 |
| **(29) Var D (%)** | 10.48 ± 1.06 | 11.11 ± 1.47 | 0.9187 | 1745 |

All variables are described by the mean ± SEM. *N*, Number of cells. < indicates significantly smaller with $p < 0.05$ ; << indicates significantly smaller with $p < 0.01$ . For most parameters, Mann–Whitney *U* test was performed between male and female mice, while fisher’s test was applied for the contingency of rebound, rebound AP, AHP and bursting.

### Castration in adult males partially de-masculinizes sexually dimorphic electrophysiological properties of mPOA neurons

To see whether sexually dimorphic electrophysiological properties in the mPOA are controlled by adult male gonadal hormones, which regulate sexually dimorphic gene expression patterns in the mPOA (Xu et al., 2012), we surveyed 72 neurons from 10 castrate male mice and 72 neurons from 9 sham controls (Figure 4A). The measured serum testosterone level confirmed successful depletion of hormones in castrates (Figure 4B, *N* = 4 sham and 5 castrates, sham, 0.52 ± 0.05 ng/ml, castrates, 0.08 ± 0.02 ng/ml, unpaired *t* test, *p*<0.0001, *t*=8.71, *df*=7). At the appearance, mPOA neurons in castrates were more female-like, smaller in the surface area and shorter in the minor axis compared to the sham control group (Table 7). Consistent with this morphological change, *C*_m_ of mPOA neurons also decreased in castrates compared to sham controls. However, for electrophysiological properties that were sexually dimorphic in intact animals, including *V*_m_, *R*_m_, *τ*_m_, AP threshold, rheobase and the percentage of rebound, only *R*_m,_ increased in castrates and became female-like while all other parameters were comparable to sham controls (Table 8). Rather, the spike half-height duration, rise time, decay time, AHP latency and Δ*G*_Sag_, which were similar in intact male and female, showed significant differences between castrate and sham controls (Table 8). Thus, depletion of adult male hormones significantly impacts electrophysiological properties of mPOA neurons but only partially de-masculinizes/feminizes sexually dimorphic electrophysiological features (Figure 5).

**Figure 4.**
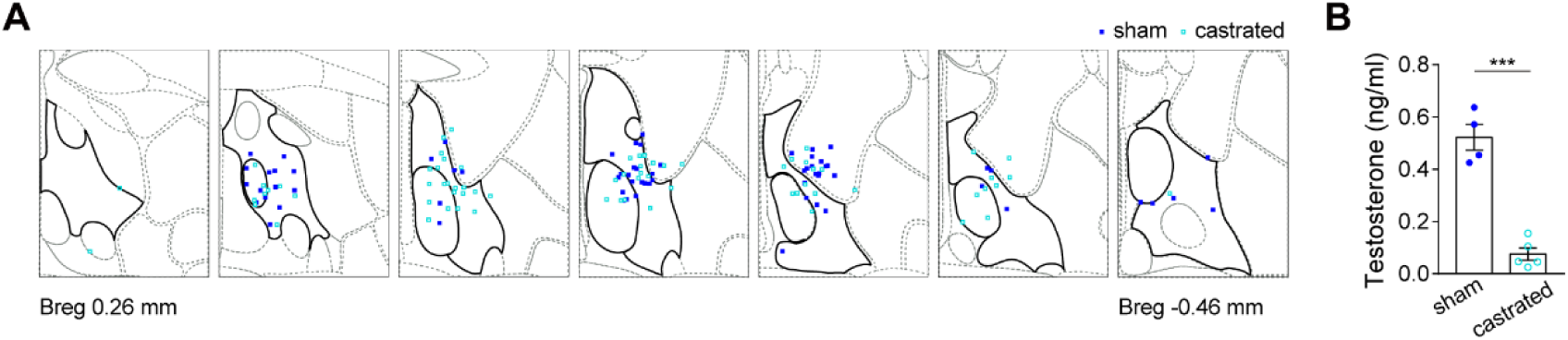
Slice recording of mPOA neurons in sham and castrated males. **A**. Positions of all neurons recorded in sham and castrated male mice, as represented by blue filled and cyan open squares respectively, were mapped onto the reference mouse brain atlas from the Allen Brain Institute (from bregma 0.26 mm to −0.46 mm). The mPOA was highlighted with thick black lines in each section. **B.** Serum testosterone level of sham and castrated male mice recorded. *N* = 4 sham and 5 castrated males. \*\*\**p* < 0.001.

**Figure 5.**
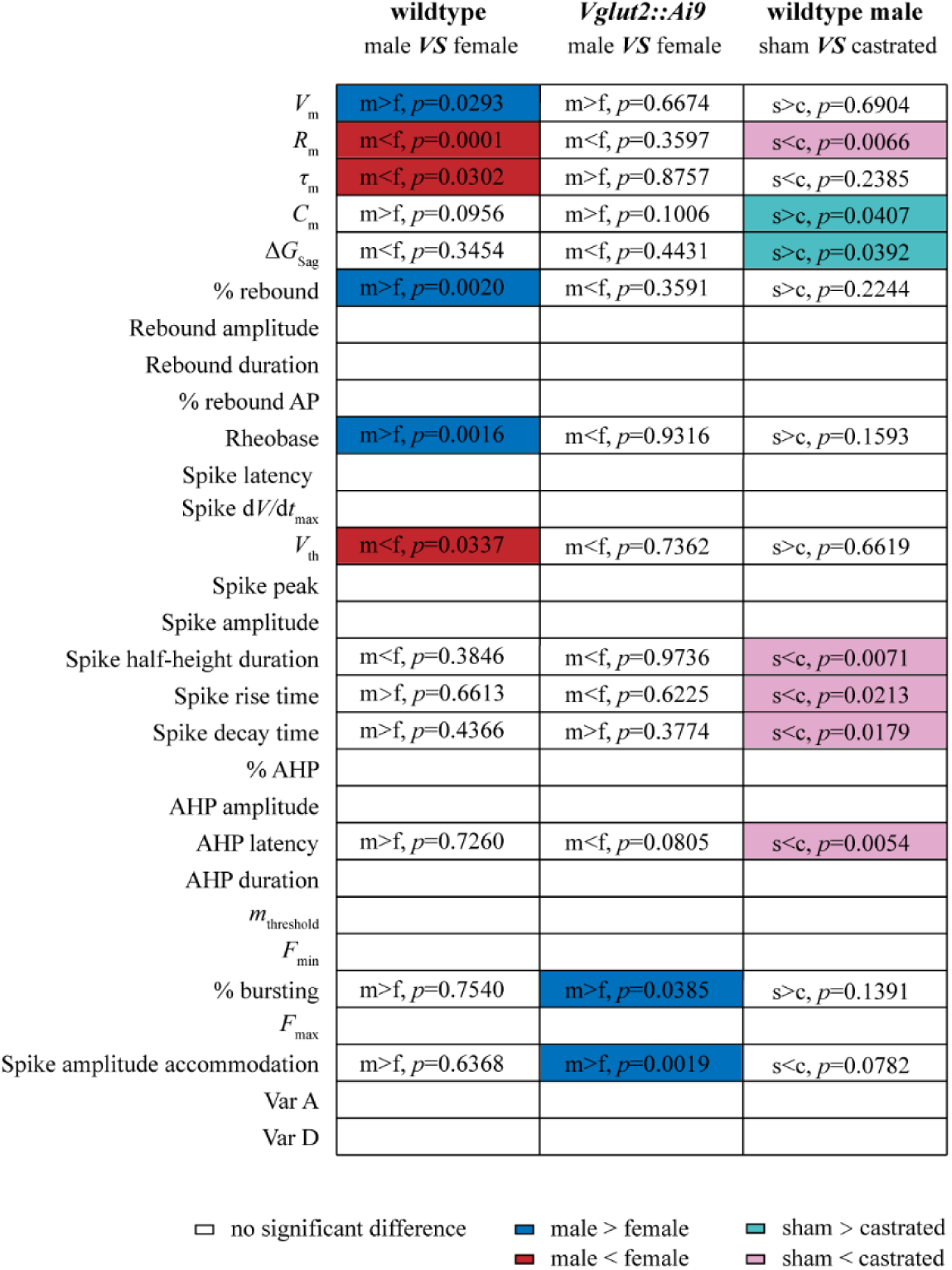
Summary of electrophysiological properties of mPOA neurons. Colored cells represent parameters that show statistically significant differences between wildtype male and female, or between *Vglut2+::Ai9* male and female, or between sham and castrated wildtype males. *p* values for parameters that show significant differences within any group were listed for all three groups. Sex differences in electrophysiological properties of mPOA neurons depend on the cell type and are only partially reversible upon castration in adult males.

**Table 7.** Soma properties of neurons in sham and castrated wildtype male mice.

|  | sham<br>(N=72) | castrated<br>(N=71) | *p* value | Mann-Whitney<br>*U* |
| --- | --- | --- | --- | --- |
| (1) soma surface<br>area (μm<sup>2</sup>) | 188.13 ± 4.58 | 171.36 ± 4.33 | 0.0088 | 1910 |
|  | castrated << sham |  |  |  |
| (2) major axis (μm) | 20.20 ± 0.36 | 19.66 ± 0.40 | 0.2600 | 2276 |
| (3) minor axis (μm) | 12.35 ± 0.22 | 11.58 ± 0.21 | 0.0080 | 1901 |
|  | castrated << sham |  |  |  |
| (4) perimeter (μm) | 56.88 ± 1.00 | 54.74 ± 0.99 | 0.1371 | 2187 |
| (5) roundness | 1.38 ± 0.03 | 1.40 ± 0.02 | 0.3109 | 2304 |

All variables are described by the mean ± SEM. *N*, Number of cells. Mann–Whitney *U* test was performed between sham and castrated male mice.

**Table 8.** Electrophysiological properties of neurons in sham and castrated wildtype male mice.

|  | sham<br>(N=72) | castrated<br>(N=72) | p value | Mann-Whitney<br>U |
| --- | --- | --- | --- | --- |
| (1) $V_m$ (mV) | -56.60 ± 0.78 | -56.69 ± 0.82 | 0.6904 | 2492 |
| (2) $R_m$ (MΩ) | 493.65 ± 28.22<br>sham << castrated | 614.06 ± 33.69 | 0.0066 | 1915 |
| (3) $\tau_m$ (ms) | 26.66 ± 1.51 | 29.15 ± 1.57 | 0.2385 | 2296 |
| (4) $C_m$ (pF) | 56.87 ± 2.41<br>castrated < sham | 49.92 ± 2.06 | 0.0407 | 2080 |
| (5) $\Delta G_{sag}$ (%) | 20.17 ± 2.22<br>castrated < sham | 19.91 ± 5.2 | 0.0392 | 2076 |
| (6) Rebound (%) | 69.44 | 58.33 | 0.2244 |  |
| (7) Rebound<br>amplitude (mV) | 4.36 ± 0.38 | 4.16 ± 0.42 | 0.5538 | 542 |
| (8) Rebound<br>duration (ms) | 274.85 ± 20.94 | 264.18 ± 21.31 | 0.7154 | 561 |
| (9) Rebound AP (%) | 18.06 | 13.89 | 0.6499 |  |
| (10) Rheobase (pA) | 23.89 ± 1.55 | 21.39 ± 1.46 | 0.1593 | 2256 |
| (11) Spike latency<br>(ms) | 132.57 ± 14.14 | 121.93 ± 12.69 | 0.3126 | 2339 |
| (12) Spike $dV/dt_{max}$<br>(mV/ms) | 327.86 ± 8.80 | 328.18 ± 11.54 | 0.9889 | 2588 |
| (13) $V_{th}$ (mV) | -34.50 ± 0.48 | -34.84 ± 0.58 | 0.6619 | 2482 |
| (14) Spike peak (mV) | 34.78 ± 0.82 | 35.93 ± 0.88 | 0.2295 | 2291 |
| (15) Spike amplitude<br>(mV) | 68.49 ± 0.94 | 70.77 ± 1.11 | 0.0626 | 2126 |
| (16) Spike<br>half-height duration<br>(ms) | 0.55 ± 0.02 | 0.61 ± 0.02 | 0.0071 | 1921 |
|  | sham << castrated |  |  |  |
| (17) Spike rise time (ms) | 0.25 ± 0.01 | 0.29 ± 0.01 | 0.0213 | 2017 |
|  | sham < castrated |  |  |  |
| (18) Spike decay time (ms) | 0.53 ± 0.05 | 0.56 ± 0.02 | 0.0179 | 2001 |
|  | sham < castrated |  |  |  |
| (19) AHP (%) | 76.39 | 63.89 | 0.1447 |  |
| (20) AHP amplitude (mV) | 15.01 ± 0.65 | 15.80 ± 0.87 | 0.3259 | 1120 |
| (21) AHP latency (ms) | 2.10 ± 0.12 | 2.77 ± 0.23 | 0.0054 | 859.5 |
|  | sham << castrated |  |  |  |
| (22) AHP duration (ms) | 4.45 ± 0.27 | 4.67 ± 0.26 | 0.2567 | 1098 |
| (23) $m_{\text{threshold}}$ (Hz/s) | -65.94 ± 56.11 | -24.90 ± 7.66 | 0.8907 | 2557 |
| (24) $F_{\text{min}}$ (Hz) | 25.68 ± 8.98 | 17.26 ± 2.06 | 0.5865 | 2455 |
| (25) Bursting (%) | 34.72 | 22.22 | 0.1391 |  |
| (26) $F_{\text{max}}$ (Hz) | 42.36 ± 3.00 | 44.24 ± 3.13 | 0.6866 | 2491 |
| (27) Spike amplitude accommodation (mV) | 10.78 ± 0.80 | 12.85 ± 0.83 | 0.0782 | 2151 |
| (28) Var A (%) | 5.98 ± 0.53 | 6.99 ± 0.93 | 0.2038 | 2273 |
| (29) Var D (%) | 8.97 ± 0.91 | 10.48 ± 1.38 | 0.9088 | 2563 |

All variables are described by the mean ± SEM. *N*, Number of cells. < indicates significantly smaller with $p < 0.05$ ; << indicates significantly smaller with $p < 0.01$ . For most parameters, Mann–Whitney *U* test was performed between sham and castrated male mice, while fisher’s test was applied for the contingency of rebound, rebound AP, AHP and bursting.

### Pharmacological blocking of T-type Ca^2+^ channel results in male mating deficits

To probe the functional relevance of the identified sex differences in electrophysiological properties of mPOA neurons, we focused on the higher percentage of rebound neurons in males after injection of hyperpolarization current. It is known that *I*_h_, mediated by hyperpolarization-activated, cyclic nucleotide-gated (HCN) channels, together with low-threshold Ca^2+^ current (*I*_T_), mediated by T-type Ca^2+^ channels, constitute currents that drive neuronal rebound (Surges et al., 2006). Given that Δ*G*_Sag,_ which is indicative of *I*_h_, is comparable between the two sexes (male, 16.81 ± 1.61%, female, 17.13 ± 2.5%, Mann–Whitney *U* test*, p*=0.3454, *U*=4345), we hypothesized that male mPOA neurons must have larger *I*_T_ to account for the higher percentage of rebound neurons. To test this more directly, we activated T-type Ca^2+^ channels by applying a −35 mV pulse (500 ms) that is preceded by a −100 mV pre-pulse (200 ms) and subtracted current recorded after application of mibefradil (20 μM), a T-type Ca^2+^ channel blocker from the current recorded before addition of the drug (Figure 6A). Indeed, the amount of Ca^2+^ entered was larger in male mPOA neurons than female (Figure 6B, male, 2.63 ± 0.43 pC, female: 1.48 ± 0.27 pC, unpaired *t* test with Welch’s correction *p*=0.0312, *t*=2.261 *df*=29.66).

**Figure 6.**
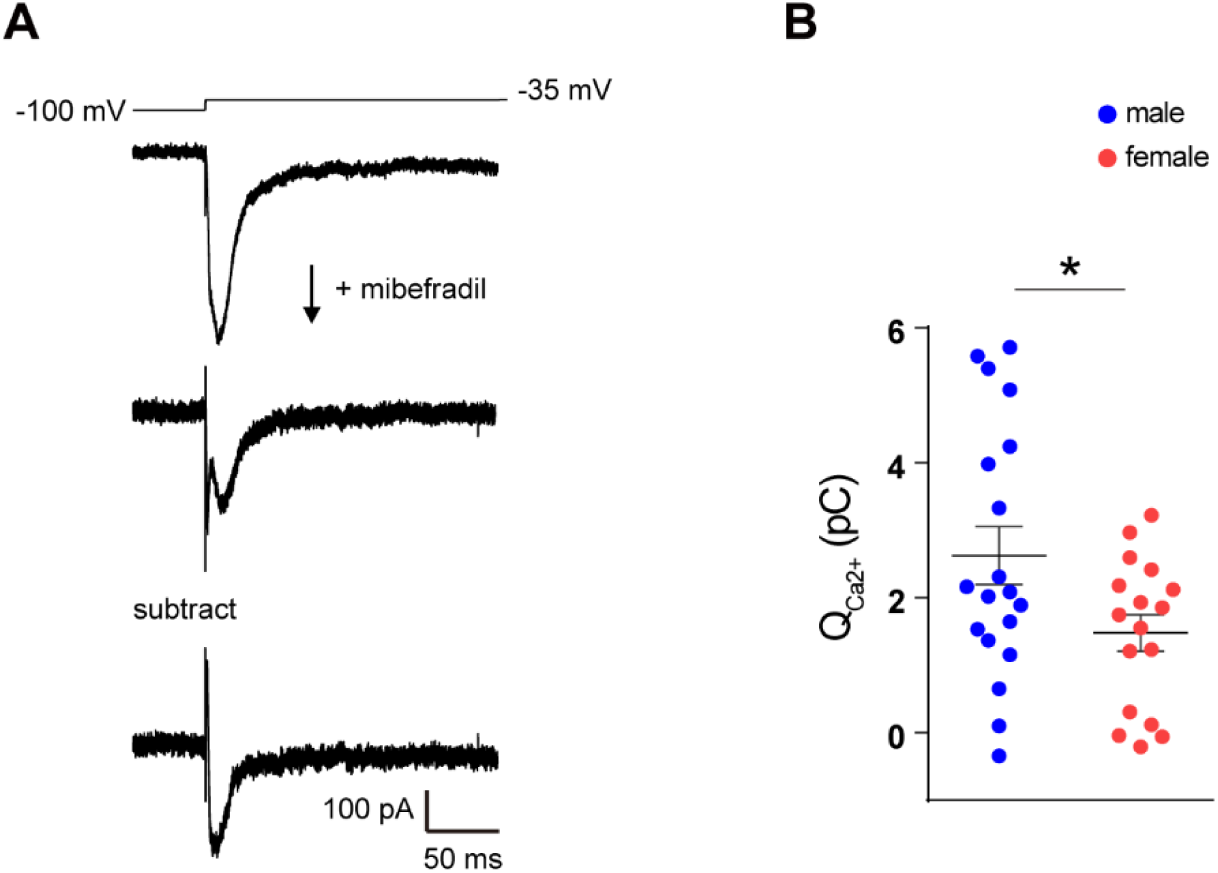
Currents mediated by T-type Ca^2+^ channels are larger in male. **A.** Representative current traces during the voltage activation from −100 mV to −35 mV (top trace) before and after applying mibefradil. The subtracted trace on the bottom illustrates Ca^2+^ currents mediated by T-type calcium channels. **B.** Quantity of Ca^2+^ entered through activated T-type calcium channels are larger in male. *N* =19 cells from 7 males and 17 cells from 6 females as represented by blue and red circles respectively. \**p* < 0.05.

Next, to test whether T-type Ca^2+^ channels play a role in male sexual behaviors we implanted bilateral cannulas into the mPOA of male mice and infused mibefradil or saline through cannulas prior to the behavioral tests (Figure 7A-B). While infusion of mibefradil did not significantly change the probability of ejaculation that occurred in the trial (Figure 7C), it nevertheless significantly decreased the number of mount and intromission without changing the count or the total duration of sniff behavior, indicating that males had difficulties to precede to consummatory acts of mating when mPOA T-type Ca^2+^ channels were blocked (Figure 7D-L). Consistent with this idea, the mean duration of sniff or mount was significantly lengthened while the total duration of intromission was decreased in mibefradil treated trials (Figure 7D-L). Thus, T-type Ca^2+^ channels appear to be important for mPOA neural dynamics and male-typical mating behaviors.

**Figure 7.**
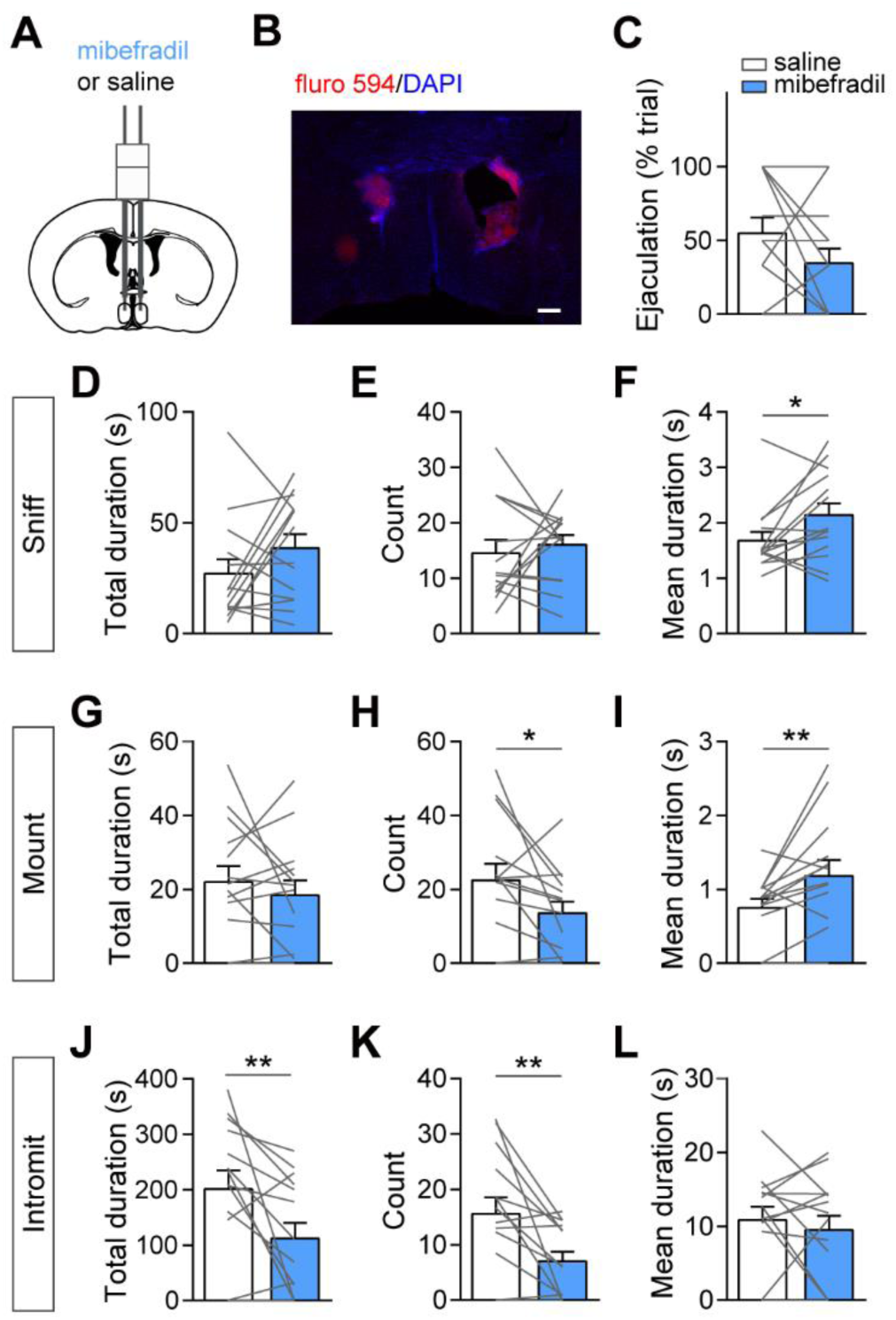
Pharmacological inhibition of T-type Ca^2+^ channels interferes with male mating. **A**. Schematics showing implantation of a bilateral cannula into the mPOA. **B.** A representative image showing *post hoc* analysis of cannula site with injection of Alexa Fluor 594. Scale bar, 200 μm. **C**. No significant effect of mibefradil on the percentage of trials that ejaculation happened. **D-F.** Mibefradil infusion did not influence the total time (D) or the number of times (E) that males sniffed the female but increased the mean duration of sniff (F). **G-I.** Mibefradil infusion did not influence the total time (G) that males mounted the female, but reduced the count (H) and increased the mean duration of mount (I). **J-L.** Mibefradil infusion reduced the total time (J) and the count (K) that males intromit a female without affecting the mean duration of intromission (L). *N* = 14. Each gray line indicates paired values from an individual animal. \**p* < 0.05; \*\**p*<0.01.

## Discussion

In this study, we compared sex differences in electrophysiological properties of mPOA neurons and investigated how sexually dimorphic features are affected by depletion of male gonadal hormones. In addition, focusing on one of the newly identified sexually dimorphic electrophysiological features, namely neuronal rebound in response to injection of hyperpolarizing current, we tested whether or how T-type Ca^2+^ channels participate in male mating behaviors. To our knowledge, this is the first time that these type of data are presented for the mPOA, an evolutionarily conserved and sexually dimorphic brain nucleus. Our study therefore exemplifies the utility of incorporating electrophysiological characterization with molecular, anatomical and behavioral studies to elucidate neural mechanisms that underlie sexually dimorphic brain functions.

Firstly, we show that mPOA neurons differ in membrane potential (*V*_m_), input resistance (*R*_m_), time constant (*τ*_m_), threshold (*V_th_*), minimum current (rheobase) required to generate an AP, and the percentage of neurons that rebound (% rebound) but not the other 23 parameters assessed. Moreover, such sex differences manifest in a cell-type dependent manner such that mPOA *Vglut2+* neurons show sex differences in the percentage of neurons that burst (% bursting) and spike amplitude accommodation but not other parameters, consistent with the known heterogeneity of the region (Tsuneoka et al., 2013, Wu et al., 2014, Tsuneoka et al., 2017, Moffitt et al., 2018, Wei et al., 2018). Importantly, there are no sex differences in either the overall composition of bursting and non-bursting cells or the gross level of excitatory of inhibitory inputs.

It is interesting that despite the pervasive sex differences in neuronal density, gene expression, synaptic organization and innervating fibers previously documented (Raisman and Field, 1973, Gorski et al., 1978, Ayoub et al., 1983, Simerly et al., 1984, Sickel and McCarthy, 2000, Xu et al., 2012), sex differences in electrophysiological properties are very restricted. These findings echo results from conventional intracellular recordings of mPOA neurons in male rodents, which have reported that different sub-regions of mPOA neurons are relative homogeneous in electrophysiological properties (Hodgkiss and Kelly, 1990, Hoffman et al., 1994). It is possible that the low throughput and *in vitro* nature of the patch-clamp or intracellular recording approach, compared to gene expression and anatomical analysis, is insufficient to capture all the distinctive sexually dimorphic features that are only present in a small fraction of mPOA neurons. Alternatively, sex differences in gene expression and other parameters may be “funneled” into subtle but functionally relevant electrophysiological properties that more directly relate to neuronal activation, neural dynamics and behaviors. Future electrophysiological characterization of more genetically defined subtypes of mPOA neurons coupled with behavioral and functional study of these populations will help to clarify the situation.

Secondly, we show that depletion of gonadal steroid hormones in adult males, while reducing neuronal size and increasing *R*_m_ such that these features become female-like in castrates, nevertheless is insufficient to de-masculinizes other sexually dimorphic electrophysiological features. Rather, castration result in significant changes in a set of parameters that are orthogonal to the panel of parameters that are sexually dimorphic in intact animals. This raise the intriguing possibility that even though both castrate males and intact females rarely display male-typical mating during behavioral tests, the underlying neurophysiological mechanisms may not be entirely the same. Thus, it may be more accurate to regard castrated males not as surgically induced “de-masculinized” or “feminized” males but rather as a peculiar state that is different from both male and female.

Finally, the higher percentage of mPOA neurons that rebound after injection of hyperpolarization current in males stood out and emerge as a possible mechanism for sexually dimorphic brain functions. It is known that *I_h_* mediated by HCN channels along with *I_T_* mediated by T-type Ca^2+^ channels drive such neuronal rebound (Surges et al., 2006). Indeed, we show that male mPOA neurons have higher *I_T_*. In fact, gonadal steroid hormones have been shown to upregulate expression of T-type Ca^2+^ channels and/or *I*_T_ in female hypothalamic neurons and in neonatal cardiomyocytes (Qiu et al., 2006, Bosch et al., 2009, Zhang et al., 2009). Importantly, pharmacological blocking of mPOA T-type Ca^2+^ channels result in difficulties for male to initiate the consummatory act of mount and to proceed into intromission after the appetitive phase of sniff behaviors. Furthermore, as T-type Ca^2+^ channels are known to be important for inhibition induced synchronized rebound firing in thalamic neurons to mediate motor functions (Anderson et al., 2005, Tai et al., 2011, Stagkourakis et al., 2018, Yang et al., 2018), it is thus tempting to speculate that T-type Ca^2+^ channels in the mPOA may participate in the sexually dimorphic ramping neural activities that is observed preceding the onset of mount and intromission (Wei et al., 2018). Future functional manipulation of specific T-type Ca^2+^ channel is required to test this exciting possibility. In summary, we have identified key sex differences in electrophysiological properties of mPOA neurons that likely contribute to sexually dimorphic display of behaviors.

## Acknowledgments

We thank members of the Xu Lab for comments on the manuscript. This work was supported by grants from the National Nature Science Foundation of China (31871066 and 31900721), Ministry of Science and Technology office China 973 program (2015CB559201), Shanghai Municipal Science and Technology Major Project (2018SHZDZX05) and the Strategic Priority Research Program of the Chinese Academy of Sciences (XDB32000000).

